# Cannabinoid receptor type 2 expression in mouse brain: from mapping to regulation in microglia under inflammatory conditions

**DOI:** 10.1101/2023.11.02.565330

**Authors:** Wanda Grabon, Anne Ruiz, Nadia Gasmi, Cyril Degletagne, Béatrice Georges, Amor Belmeguenai, Jacques Bodennec, Sylvain Rheims, Guillaume Marcy, Laurent Bezin

## Abstract

Since its detection in the brain, the cannabinoid receptor type 2 (CB2) has been deemed a promising therapeutic target for various neurological and psychiatric disorders. However, precise brain mapping of CB2 expression is currently lacking. Using magnetic cell sorting, calibrated reverse transcription-quantitative PCR, and single-nucleus RNA-seq, we revealed the low level of CB2 expression in all brain regions examined, mainly by a few microglial cells and by neurons in an even lower proportion. Upon lipopolysaccharide stimulation to simulate non-sterile neuroinflammatory conditions, we demonstrated that the inflammatory response was associated with a transient reduction in CB2 mRNA levels in the brain tissue, particularly in microglia. This result, confirmed in the BV2 microglial cells, contrasts with the positive correlation observed between CB2 mRNA levels and the inflammatory response upon stimulation by interferon-gamma, which models sterile inflammatory conditions. Thus, discrete brain CB2 expression may be upregulated or downregulated depending on the inflammatory context.

## 1. INTRODUCTION

The function of cannabinoid receptor type 2 (CB2), mainly expressed by leukocytes, was initially limited to its peripheral immunomodulatory role ^1–3^. However, the use of CB2-specific ligands and the availability of CB2-knockout (KO) mice have revealed their potential role in the central nervous system (CNS) functions under both physiological and pathological conditions ^4,5^. To gain deeper insights into the involvement of CB2 in brain function, it is essential to identify cells targeted by CB2 ligands, which hold substantial therapeutic promise in neuropsychiatric and neuroinflammatory diseases ^6,7^. Nevertheless, precise and accurate mapping of *Cnr2* gene expression remains difficult to establish based on CB2 protein detection, primarily because specific antibodies are still lacking ^8,9^; therefore, this can only be achieved by CB2 transcript detection.

The primary goal of the current study was to quantify and compare tissue expression levels of CB2 and other primary cannabinoid receptors, i.e., CB1, GPR18, and GPR55, across six major brain regions in three different mouse strains (C57bl/6, Balb/c, and Swiss) using calibrated reverse transcription-quantitative PCR (RT-qPCR). The second objective was to determine cell types that predominantly support CB2 expression in the brain. We used cell-sorting techniques on dissociated adult brain tissue to determine CB2 mRNA levels in populations enriched in microglia and neurons, astrocytes, oligodendrocytes, and endothelial cells. To achieve this objective, we took a particular attention in limiting microglial cell activation across the entire technical procedure using transcriptional and translational inhibitors ^10^. Furthermore, we accessed a single-nucleus database derived from the cortical tissue collected from 10-day-old mice to assess the proportion of CB2-expressing cells in each cell population.

CB2 expression is known to be regulated by inflammation ^11,12^. Indeed, CB2 expression in brain tissue was found to be induced in neurological conditions associated with an inflammatory state, such as stroke, traumatic brain injury, and Alzheimer’s disease ^5,7^. Most studies suggest that microglial activation is responsible for CB2 induction. However, a few *in vitro* studies that simulated non-sterile neuroinflammation by stimulating microglial cells with lipopolysaccharide (LPS) reported the downregulated expression of CB2 ^12,13^. Our third objective was to comprehensively clarify whether CB2 expression coordinated with that of inflammatory markers after a sterile or non-sterile inflammatory challenge, modeled by stimulation with interferon-gamma (IFNγ) or LPS, respectively, which are known to activate different inflammatory intracellular signaling pathways ^14,15^. To the best of our knowledge, this study is the first to demonstrate that CB2 gene expression in the brain is supported by a very small number of microglial cells and by neurons in an even smaller proportion and that microglial CB2 transcript levels can be up- or downregulated depending on the inflammatory context.

## 2. METHODS

Full details of the methods are given in the supplementary material.

### Experimental design

**Experiment 1**. To assess mRNA level of CB2, CB1, GPR55 and GPR18 in different mouse strains and brain regions, brains from adult male Balb/c, C57Bl/6 and Swiss mice (n=3/group) were collected after transcardiac perfusion of saline, and olfactory bulbs, neocortices, hippocampi, hypothalamus, cerebellum and brainstem were microdissected. Transcript levels were determined by RT-qPCR.

**Experiment 2**. To validate the efficacy of transcription and translation inhibitors in limiting *ex vivo* microglial cell activation induced during tissue dissociation and magnetic sorting, the protocol was run in parallel using 3 different buffer conditions applied to all steps of the protocol, from intracardiac perfusion to collection of sorted cells. Experiment was performed on adult C57Bl/6 male mice (n=2/buffer condition). Transcript levels of inflammatory genes were determined by RT-qPCR.

**Experiment 3**. To determine in which brain cell types CB2 is expressed under physiological state, brains of C57Bl/6 mice were collected after transcardiac perfusion of the buffer selected in experiment 2 (n=3). Neocortices and hippocampi were microdissected and immediately processed for magnetic cell separation. Transcript levels were determined by RT-qPCR. Furthermore, a single-nucleus database obtained from cortex tissue collected from 10-day-old mice was used to estimate the proportion of CB2-expressing cells in each cell population.

**Experiment 4**. To measure the effect of inflammation on CB2 expression in brain microglia, adult C57Bl/6 mice were treated intra-peritoneally (IP) with LPS (*Escherichia coli* O55:B5, Sigma, L2880) at 5 mg/kg and brains were collected 3h (n=5) or 24h (n=5) later after transcardiac perfusion of saline. Mice treated with 0.9% NaCl were used as controls (n=4). Neocortices and hippocampi were microdissected. Transcript levels were determined by RT-qPCR.

**Experiment 5**. To investigate CB2 expression in microglial cells under physiological and inflammatory condition, murine BV2 cells were cultured with or without LPS (100 ng/mL) or IFNγ (100 ng/mL) and harvested 1h-24h later for further RT-qPCR analysis (CTRL, LPS+1h, +2h, +4h: 4 wells/condition; LPS+8h, +24h: 3 wells/condition; CTRL, IFNγ+3h, +8h: 3 wells/condition). To determine the role of IL-1R signaling in regulating CB2 expression, BV2 cells incubated with LPS (100 ng/mL) were pre-treated for 30 min and co-cultured for 2h with IL1-RA, before harvesting and further RT-qPCR analysis.

### Animals

Adult male mice (Balb/c, C57Bl/6 and Swiss, 8-weeks-old, Envigo, France) were used in this study. The experimental procedures were conducted in accordance with the European Community guidelines for care in animal research and approved by the CELYNE local Ethics Research Committee (protocol #24302). Every effort was made to minimize animal suffering.

### BV2 cell culture

The immortalized murine BV2 cell line (BV2 cells) was kindly provided by Dr. Nadia Soussi (NeuroDiderot, Paris University). Cells at passage 9 to 15 were treated with LPS (from *Escherichia coli* O55:B5, 100 ng/mL, Sigma #L6529) or with IFNγ (100 ng/mL, Gibco #PMC4031) for the indicated time, and harvested for further RT-qPCR analysis. Blockade of IL-1R was performed using the recombinant mouse IL1-RA protein (1-1000 mg/mL, abcam #ab283475).

### Reverse transcription and real-time quantitative PCR

After extraction, total RNAs were reverse transcribed to complementary DNA (cDNA) using both oligo dT and random primers with PrimeScript RT Reagent Kit (Takara, #RR037A) according to manufacturer’s instructions in the presence of a synthetic external non-homologous poly(A) standard messenger RNA (SmRNA; A. Morales and L. Bezin, patent WO2004.092414) to normalize the RT step, as previously described ^16^. Each cDNA of interest was amplified using the Rotor-Gene Q thermocycler (Qiagen), the SYBR Green PCR kit (Qiagen, #208052) and oligonucleotide primers (Eurogentec) specific to the targeted cDNA. cDNA copy number detected was determined using a calibration curve, and results were expressed as cDNA copy number/μg tot RNA.

Pro-inflammatory index was calculated using a specific set of pro-inflammatory genes: IL-1β, IL-6 and TNFα. For each sample, the number of copies of each transcript has been expressed in percent of the averaged number of copies measured in the whole considered group of samples. Once each transcript was expressed in percent, an index was calculated by adding the percent of each transcript involved in the composition of the index and expressed in arbitrary units (A.U.).

### Brain dissociation and magnetic cell sorting (MACS studies)

To prevent any artifactual *ex vivo* gene expression changes during brain dissociation and cell sorting procedures, all buffers and solutions used during the process (from animal perfusion to sorted cells flash freezing) were supplemented with a cocktail composed of Actinomycin D (3μM, Tocris #1229/10), Anisomycin (100 μM, Tocris #1290/50) and Triptolide (10 μM, Tocris #3253/10)^10^. All steps were performed on ice or using pre-chilled refrigerated centrifuge set to 4°C with all buffers/solutions pre-chilled before addition to samples to further limit cell activation. The general workflow of brain dissociation and magnetic cell sorting is illustrated in supplementary data (**Fig. S1**).

### Brain dissociation and Magnetic Cell-sorting

Neocortices and hippocampi were quickly dissected, cut in smaller pieces and processed for dissociation using Miltenyi’s Adult Brain Dissociation Kit (#130-107-677) according to manufacturer’s instruction. To enhance cell yields, distinct mice were used for the isolation of neurons (n=3) and for the isolation of the other cell types (microglia, endothelial cells and astrocytes, n=3). Neurons were enriched using Adult Neuron Isolation Kit (Miltenyi #130-126-602). Endothelial cells, microglia and astrocytes were magnetically sorted successively, using the anti-Ly-6C biotin antibody (Miltenyi #130-111-914) and Anti-Biotin MicroBeads (Miltenyi #130-090-485), the CD11b MicroBeads (Miltenyi #130-093-634), and the ACSA-2 MicroBeads (Miltenyi #130-097-679), respectively. Sorted cells were counted manually, spun and dry cell pellets were flash frozen and stored at −80°C.

### Flow Cytometry

To control purity of enriched cell populations, a fraction of cell suspensions was collected before and after each sorting. The following antibodies were added to cell suspensions at 1:50 concentration in B2 buffer: ACSA2-APC (Miltenyi #130-116-245), CD11b-PE Vio770 (Miltenyi #130-113-808), CD31-PE (Miltenyi #130-111-540), Ly6C-APC Vio770 (Miltenyi #130-111-919) and O4-VioBright515 (Miltenyi #130-120-102). DAPI (2.5 μg/mL, Sigma #D9542) was added as a viability marker right before flow cytometry analysis. Cells were analyzed with the BD FACS Canto II Flow Cytometer (Becton-Dickinson) and data files with FlowJo software V10.7.2 (Becton-Dickinson).

### Brain dissociation and single nucleus RNAseq (single nucleus study)

#### Tissue dissociation

Nuclei from whole cortex were obtained from one mouse at age P10, anesthetized with isoflurane and sacrificed by decapitation. The dissected cortex tissue was immediately placed in a dry-ice-cold tube for immediate freezing until processing for nuclei isolation.

#### Single-nucleus isolation

Dissected frozen cortex was resuspended and mechanically homogenized using dounce homogeneizer to release nuclei following the Salty EZ 10 protocol (dx.doi.org/10.17504/protocols.io.bx64prgw). Dissected frozen cortex was resuspended into 600μL of cold homogenization buffer that consisted of 10mM Tris HCl pH 7.5, 146mM NaCl, 1mM CaCl_2_, 21mM MgCl_2_, 0.03% Tween20, 0.01% BSA, 10% EZ buffer (Sigma) and 0.2U/μL Protector RNase Inhibitor (Roche). Tissues were then transferred into 2mL dounce (Kimble) and homogenized using 10 strokes of the loose pestle followed by 8 strokes of the tight pestle to release nuclei, on ice. Homogenate was then strained through a 70μm cell strainer (Pluriselect) and centrifuged at 500g for 5 minutes to pellet nuclei. After removing supernatant, nuclei were washed in 1ml resuspension buffer containing 10mM Tris HCl pH 7.5, 10mM NaCl, 3mM MgCl_2_, 1% BSA and 0.2U/μL Protector RNase Inhibitor and centrifuged at 500xg for 5 minutes. Nuclei were then resuspended in 500μL of resuspension buffer and 5.10^5^ of the best singlet nuclei were sorted (BD ARIA) based on DAPI intensity before counting using the LUNA automated cell counter. Nuclei were finally centrifuged at 500xg for 5 minutes and diluted in resuspension buffer to a concentration of 1200 nuclei/μl before encapsulation in 10x Chromium. All steps were carried on ice or at 4°C.

#### Single-nucleus capture and sequencing

Single-nuclei capture and sequencing were performed at the Cancer Genomics Platform of the Cancer Research Center of Lyon (CRCL). Nuclei suspension (1,200 nuclei/μL) were loaded onto a Chromium iX (10X Genomics) to capture 10,000 single nuclei. cDNA synthesis and library preparation were done following the manufacturer’s instructions (chemistry V3.1) and library has been sequenced using the Novaseq 6000 (Illumina) to reach 30k reads per nucleus. Cell Ranger version 6.1.1 (10X Genomics) was used to align reads on the mouse reference genome gex-mm10-2020-A and to produce the count matrix.

#### Single-nucleus RNA-seq data analysis

The gene expression matrices from Cell Ranger were used for downstream analysis using the software R (version 4.1.2) and the R toolkit Seurat (version 4.1.0). Nuclei were excluded from downstream analysis when they had more than 3% mitochondrial genes, fewer than 300 unique genes, more than 20,000 unique molecular identifiers (UMIs) and detected as doublets using scdblFinder R package. A total of 8,530 cells were selected. Gene expression was normalized using the standard Seurat workflow and the 2,000 most variable genes were identified and used for principal component analysis (PCA). The top most significant principal components (PCs) were selected for generating the UMAP, based on the ElbowPlot method in Seurat. Clustering of cells was obtained following Seurat graph-based clustering approach with the default Louvain algorithm for community detection. We then performed differential expression analysis using the FindMarkers function of Seurat with the default Wilcoxon rank sum test and annotated clusters based on expression of marker genes (**Fig. S4**). We then manually annotated the major classes of cells: Neurons, Microglia, Astrocytes, Oligodendrocytes and vascular cells (**Fig. 3H-I**).

### Statistical Analysis

Statistical analyses were performed using Prism 10.0 software (GraphPad, USA). Results are presented as mean ± SEM (standard error of the mean). Differences with a p-value<0.05 (p<0.05) were considered to be statistically significant. The Shapiro–Wilk test and quantile–quantile plot were used to assess normal distribution of the data. For normal data, the statistical significance was assessed by two-tailed t-test or one-way ANOVA, followed with Tukey’s *post-hoc* test for multiple comparisons. For non-normal data, the statistical significance was assessed by Kruskal-Wallis test, followed with Dunn’s *post-hoc* test for multiple comparisons.

## 3. RESULTS

### 3.1. CB2 expression in basal condition is consistent across brain regions with minimal strain variability

Tissue transcript levels of CB2 and three other cannabinoid receptors (CB1, GPR18, and GPR55) were determined in six microdissected regions of the adult (8-week-old) mouse brain: olfactory bulb, neocortex, hippocampus, hypothalamus, cerebellum, and brainstem (**Fig. 1A**). A comparative analysis of cannabinoid receptor mRNA levels was performed across three commonly used mouse strains: C57bl/6, Balb/c, and Swiss (n=3/group). Transcripts of the four receptors were detected in all mouse strains and examined brain regions. From the 72 statistical comparisons (3 strains, 6 brain structures, 4 receptors), a single difference was observed between Balb/c and C57bl/6 mouse strains for GPR18 in the neocortex (Tukey’s multiple comparison test following one-way ANOVA: p=0.0338; **Table S2**). Therefore, to compare the expression of CB2 with that of the other cannabinoid receptors in different regions, data from the three mouse strains were combined (n=9). For CB2 transcripts, no significant inter-region difference (p=0.1801) was observed, with an average of 1,112 ± 73 copies/μg RNA (**Fig. 1B**).

**Figure 1.**
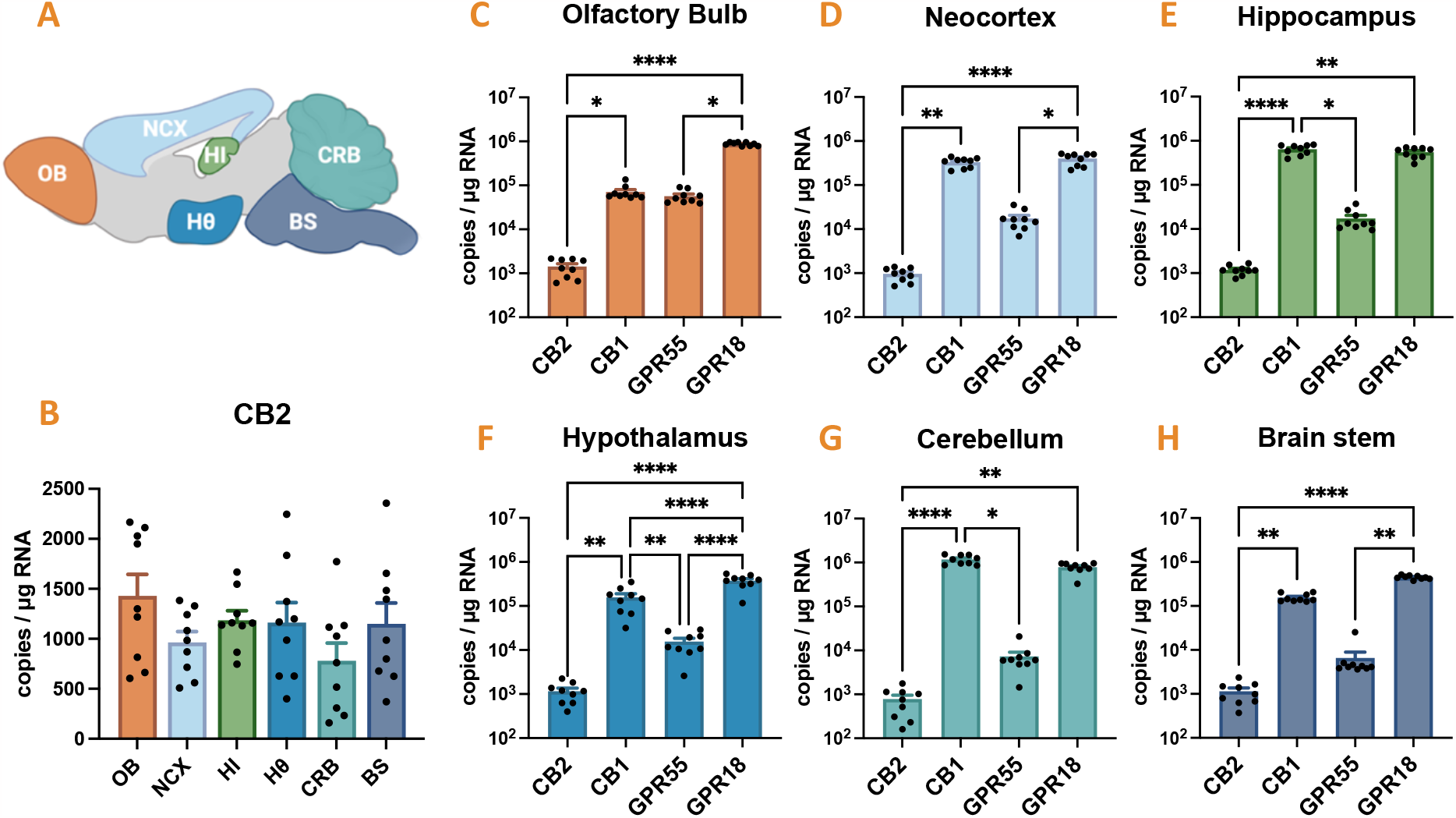
Regional distribution of cannabinoid receptors mRNA in the healthy mouse brain. CB2 mRNA was quantified using calibrated RT-qPCR and compared with other cannabinoid receptors CB1, GPR18 and GPR55 in six microdissected regions from adult mouse brain: the olfactory bulb (OB), the neocortex (NCX), the hippocampus (HI), the hypothalamus (Hθ), the cerebellum (CRB) and the brainstem (BS), n=9 mice. **A**. Representation of the 6 microdissected regions. **B**. Quantification of CB2 transcript level. One-way ANOVA: p=0.1801. **C-H**. Quantification of CB2, CB1, GPR18 and GPR55 in each microdissected region. Transcript levels are expressed as copies of cDNA per microgram of total RNA. Values are presented as means + SEM. Normal data were analyzed with Tukey’s post-hoc test for multiple comparisons following one-way ANOVA (Hθ: p<0.0001). Non-normal data were analyzed with Dunn’s post-hoc test for multiple comparisons following Kruskal-Wallis test (OB: p<0.0001, NCX: p<0.0001, HI: p<0.0001, CRB: p<0.0001, BS: p<0.0001). *, p<0.05; **, p<0.01; ***, p<0.001, ****, p<0.0001. Not significantly statistical differences are not graphically represented.

It should be noted that the distribution of transcript levels of the other cannabinoid receptors was markedly less homogenous (Kruskal-Wallis: CB1, p<0.0001; GPR18, p<0.0001; GPR55, p<0.0001; **Fig. S2**). In all examined regions, CB2 expression was lower than that of other endocannabinoid receptors (**Fig. 1C-H**). Overall, CB1 and GPR18 were the two most highly expressed cannabinoid receptors among examined regions, with GPR55 expressed at intermediate levels.

### 3.2. Transcription and translation inhibitors prevent microglia *ex vivo* activation during tissue dissociation and cell sorting

*Ex vivo* activation of brain cells, particularly microglial cells, can alter the accuracy of measurements related to the inflammatory state within the brain tissue. Although this activation can be minimized by rapid microdissection on ice and immediate freezing of samples, it remains a concern during tissue dissociation protocols involving heating, enzymes, mechanical damage, and cell sorting ^10,17^. To prevent this activation, buffers can be supplemented with transcriptional and translational inhibitors, such as actinomycin D (ActD), triptolide (Trip), and anisomycin (Anis) ^17^. Herein, we examined the efficacy of these inhibitors in CD11b-enriched populations from adult C57bl/6 mouse brains, with purity reaching 91.1% (**Fig. 2A**). Based on quantification of *IL-1β* and *TNFα* transcripts, the level of gene activation in microglial cells sorted from adult mouse brains was the highest when the buffer contained no inhibitor (**Fig. 2B-C**). For the *IL-1β* transcript, ActD-induced inhibition of transcription alone decreased its expression level by 1.9-fold, whereas inhibition of both transcription and translation using the inhibitor cocktail decreased *IL-1β* expression by 11.9-fold (**Fig. 2B**). For the *TNFα* transcript, decreased expression was only observed with the application of the inhibitor cocktail (**Fig. 2C**). Our data suggest that the inhibition of both transcription and translation, applied as early as intracardiac perfusion, is crucial to limit gene activation in microglial cells during tissue dissociation and cell sorting ^17^. Therefore, to avoid any bias related to the *ex vivo* activation of microglia, all further magnetic-activated cell-sorting (MACS) studies were performed in the presence of an inhibitor cocktail for brain perfusion, dissociation, cell labeling, and cell sorting.

**Figure 2.**
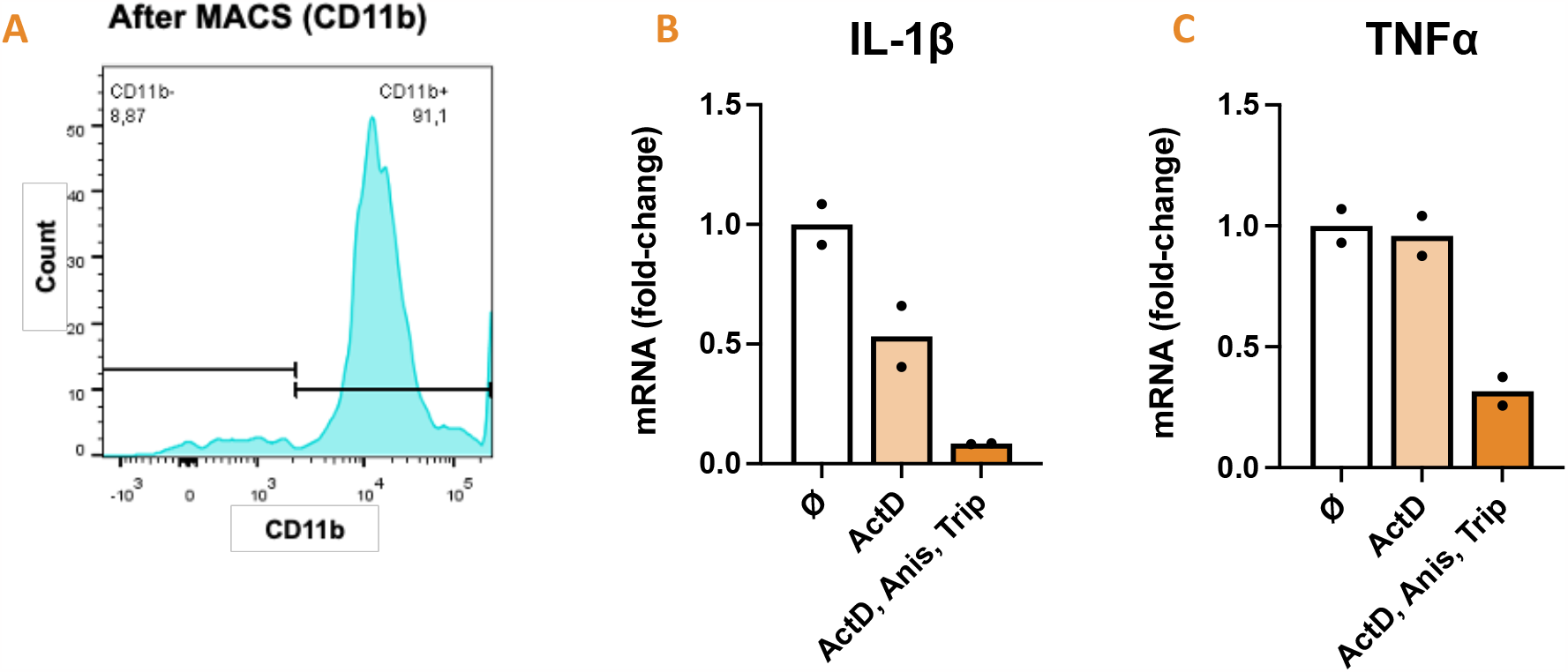
Prevention of *ex vivo* microglial activation during tissue dissociation and cell sorting protocols using transcription and translation inhibitors. **A**. Brain tissue from adult mice was dissociated using Miltenyi’s ABDK protocol and CD11b-positive cells were enriched using MACS CD11b magnetic microbeads. Purity of CD11b-positive population was controlled on a pool of cells from each sample by cytometry. **B-C**. IL-1β (B) and TNFα (C) mRNAs were quantified using calibrated RT-qPCR following tissue dissociation and CD11b MACS cell enrichment from adult mouse brain (n=2). Dissociation and MACS protocols were performed with classic buffer (ø), with buffer complemented with transcription inhibitor ActD (3 μM) or with buffer complemented with both transcription and translation inhibitors ActD, Anis and Trip (3 μM, 100 μM and 10 μM, respectively). Transcript levels are expressed as fold change of the classic buffer (ø) condition. Individual values (dots) and average values (bars) are presented.

### 3.3. CB2 mRNA expression in sorted brain cells shows peak levels in microglia at the physiological state

Distinct cell populations from the mouse hippocampus and neocortex were enriched using MACS after tissue dissociation and subsequently immunophenotyped using flow cytometry (**Fig. S3**). High degrees of purity were observed, with 98.2% purity in cell population enriched in both astrocytes (ACSA2^+^/O4^-^: 64%) and oligodendrocytes (O4^+^: 34.2%), 60.9% purity in a cell population enriched in Ly6C^+^/CD31^+^ endothelial cells, 98.2% purity in cell population enriched in CD11b^+^ microglial cells, and 96.9% purity in the neuronal population negative for all tested flow cytometry cell markers. The identity of the enriched cell populations was further confirmed by RT-qPCR by detecting and quantifying the transcript levels of various cell markers (**Fig. 3A-D**). Notably, CB2 mRNA levels differed significantly between distinct enriched cell populations (**Fig. 3E**). The highest CB2 mRNA expression was found in the microglia-enriched cell population, showing a 10-fold increase compared with the neuronal cell-enriched population and a 63- to 153-fold increase compared with endothelial cells and astrocytes/oligodendrocytes, respectively.

**Figure 3.**
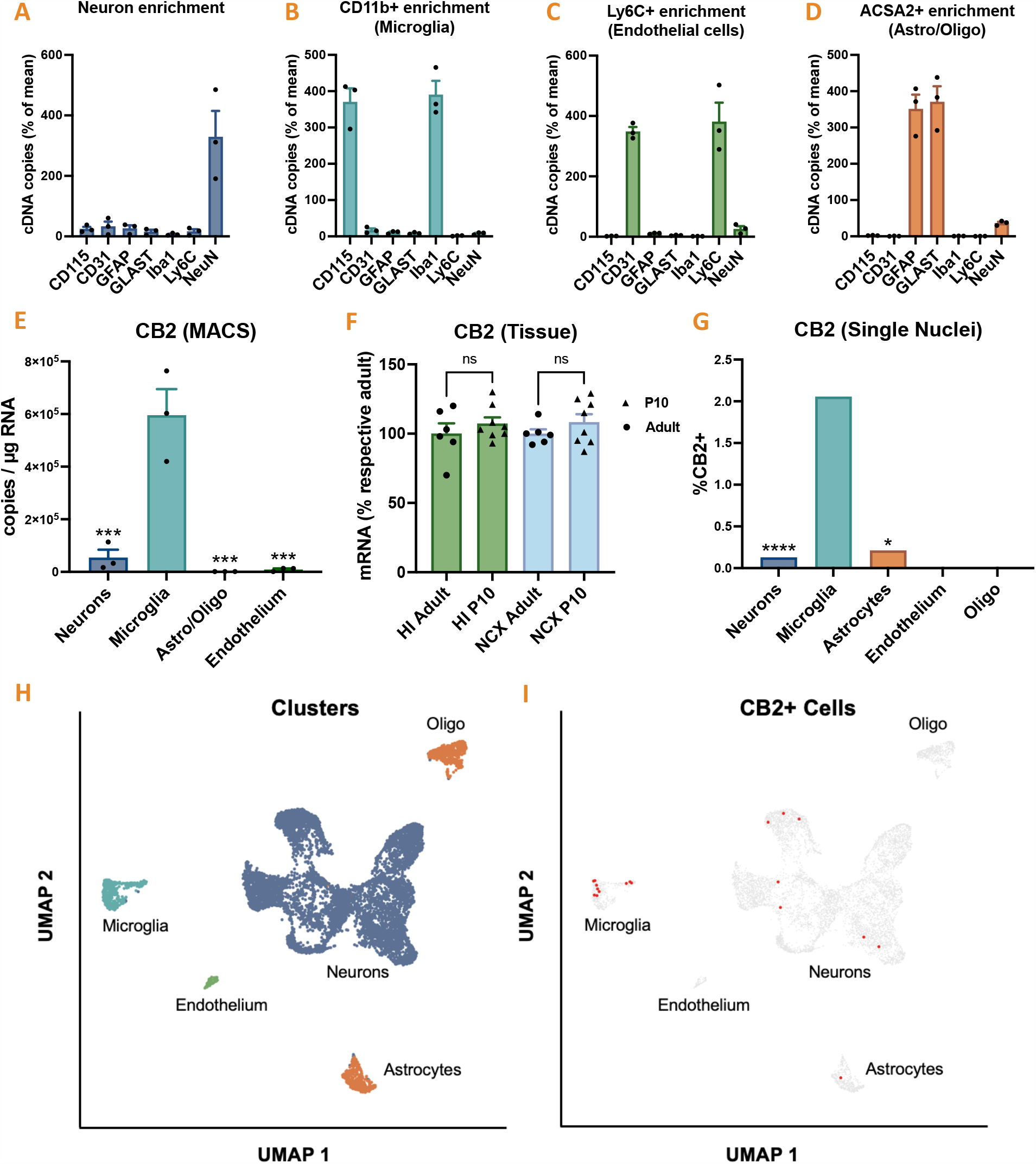
Cell distribution of CB2 mRNA in the mouse brain. **A-E**. Adult C57Bl/6 mouse brain tissue was dissociated as in Fig. 2. Four populations were enriched by MACS technique. Expression of specific cell markers was quantified by RT-qPCR to control for cell population identity and purity: **A**. Neurons, expressing NeuN mRNA; **B**. CD11b-positive microglia, expressing CD115 and Iba1 mRNA; **C**. ACSA2-positive astrocytes and oligodendrocytes, expressing GFAP and GLAST mRNA; **D**. Ly6C-positive endothelial cells, expressing CD31 and Ly6C mRNA. **E**. CB2 mRNA was quantified using calibrated RT-qPCR in the four MACS-enriched cell populations. Transcript levels are expressed as copies of cDNA per μg of total RNA reverse transcribed. Values are presented as mean + SEM, analyzed with Tukey’s test for multiple comparisons following one-way ANOVA. 554,052 ± 92,463 CB2 cDNA copies/μg RNA were quantified in microglia-enriched cell population. Microglia vs. Neurons: p=0.0004; Microglia vs. Astro/oligo: p=0.0002; Microglia vs. Endothelial cells: p=0.0002. **F**. Levels of CB2 transcripts quantified at tissue level in the hippocampus (HI) and neocortex (NCX) are expressed as % of the mean calculated in respective adult group. CB2 mRNA levels are similar in P10 (n=8) and adult (P56) mice (n=6), analyzed with Tukey’s test for multiple comparisons following one-way ANOVA. HI adult vs. P10: p=0.7796; NCX adult vs. P10: p=0.6922. **G**. Percentage of CB2-positive cells in clustered nuclei. Data are analyzed with Fisher exact test. Microglia vs. neurons, p<0.0001; microglia vs. astrocytes: p=0.0114. **H**. Clustering of Single nuclei sorted from P10 mice neocortices **I**. Visualization of CB2-expressing nuclei (red) among clustered sorted nuclei. Asterisks represent comparisons vs. microglia: *, p<0.05; **, p<0.01; ***, p<0.001, ****, p<0.0001.

Next, we performed single-nucleus RNA-seq on 10-day-old mice to estimate the proportion of CB2-expressing cells in each population enriched by MACS. Neonatal brain tissue has the advantage of being easy to dissociate for subsequent single-cell analysis, unlike adult tissue, which contains large amounts of debris due to myelin accumulation. Importantly, the stability of CB2 mRNA levels between the neonatal brain and adult brain in both the hippocampus and neocortex (**Fig. 3F**) facilitated a meaningful comparison between the MACS data obtained at 8 weeks and the single-nucleus RNA-seq data obtained at 10 days.

We isolated 10k nuclei, excluded low-quality nuclei, and clustered and annotated cell types (**Fig. 3H**) based on classical markers (**Fig. S4**). CB2 mRNA was detected in less than 1% of all cells, identified as neurons, microglia, and astrocytes. Microglial cells had the highest proportion of CB2-positive cells, approximately 20-fold greater than that observed in neurons and 10-fold greater than that in astrocytes. The CB2-expressing nuclei are shown in red in **Fig. 3I**.

### 3.4. CB2 mRNA levels in the neocortex and the hippocampus following LPS

In adult C57bl/6 mice, LPS administration led to a significant inflammatory peak after 3 h, as reflected by high levels of *TNFα*, IL-*1β, IL-6, COX2, NOS2*, and *MCP1* transcripts in the hippocampus and neocortex (**Fig. 4A-F**). The inflammatory index calculated from the transcript levels of these pro-inflammatory genes peaked at 13.2-fold ± 1.6 of the control level (**Fig. 4G**). Three hours after LPS administration, i.e., during the inflammatory peak, the CB2 transcript level transiently decreased to 52 ± 9% of that in controls, subsequently rebounding 24 h later to 194 ± 19% above controls once inflammation resolved (**Fig. 4H**). The CB2 transcript level was inversely correlated with the inflammatory index after LPS administration (**Fig. 4I**). Conversely, there was no association between CB1 and GPR18 transcript levels, whereas GPR55 transcript levels positively correlated with the inflammatory index (**Fig. S5**).

**Figure 4.**
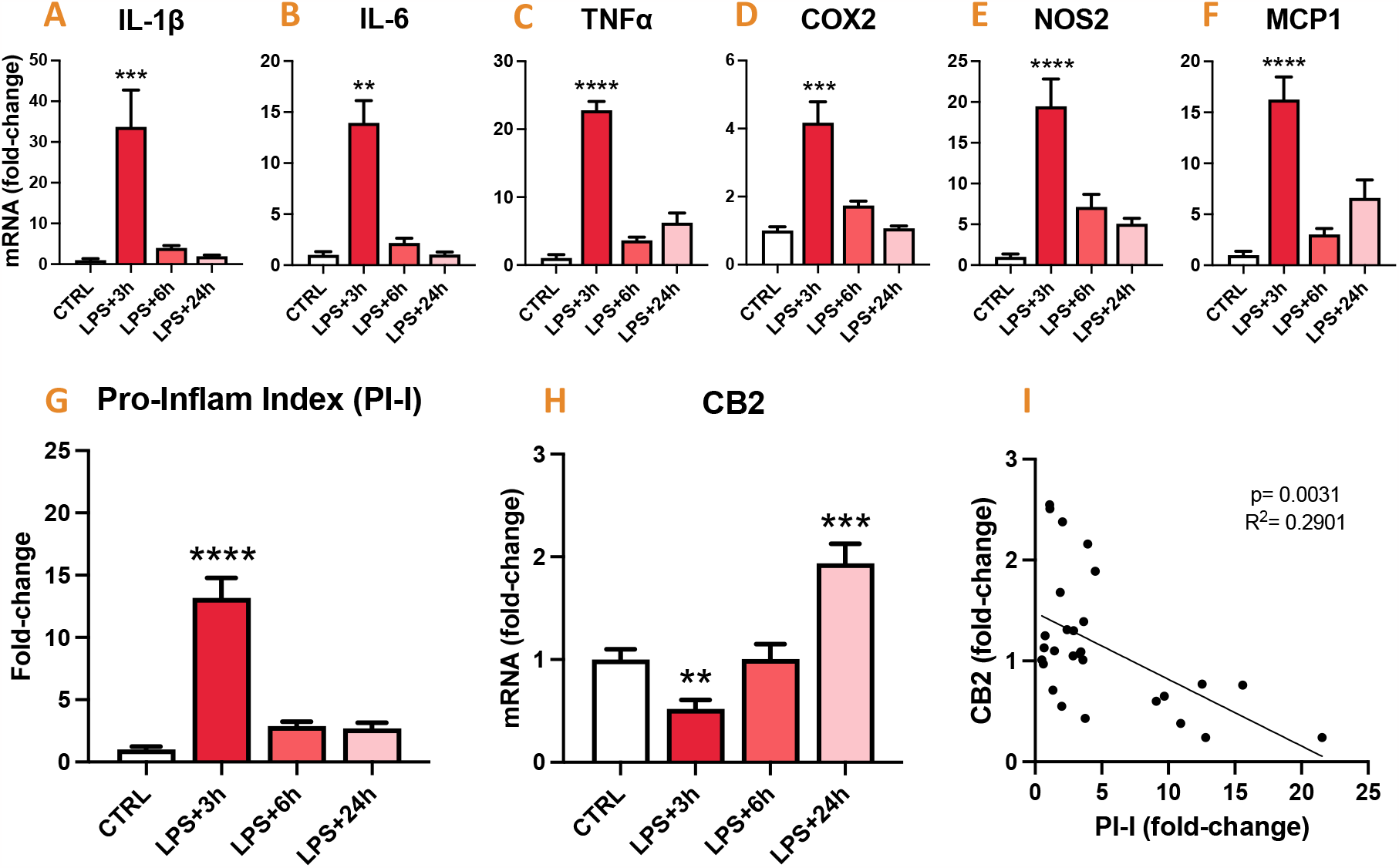
Tissue expression of CB2 mRNA inversely correlated with that of inflammatory markers in the mouse brain after LPS challenge. Transcript levels of pro-inflammatory genes and CB2 were quantified in the neocortex and the hippocampus of adult C57Bl/6 mouse brains, 3h (n=7), 6h (n=7) and 24h (n=8) following LPS administration (5 mg/kg, IP) and compared with untreated control (CTRL) mice (n=6). **A-F**. Quantification of IL-1β, IL-6, TNFα, COX2, NOS2 and MCP1 transcript levels. **G**. Pro-inflammatory index (PI-I) was calculated from transcript levels of pro-inflammatory genes (A-F). CTRL vs. LPS+3h: p<0.0001. **H**. Quantification of CB2 mRNA. Data are expressed as the mean ± SEM, and presented as relative to control mice. CTRL vs. LPS+3h: p=0.0036; CTRL vs. LPS+24h: p=0.0008. Normal data (NOS2, MCP1, PI-I and CB2) were analyzed with Tukey’s test following one-way ANOVA. Non-normal data (IL-1β, TNFα, IL-6, COX2) were analyzed with Dunn’s test following Kruskal-Wallis test. Asterisks represent LPS-treated groups vs. CTRL: *, p<0.05; **, p<0.01; ***, p<0.001, ****, p<0.0001. **I**. Relationship between CB2 transcript level and PI-I (simple linear regression, p=0.0031, R2=0.2901, y=-0.06617x+1.480, n=28).

### 3.5. CB2 mRNA levels in microglia following LPS

All steps were performed using buffers complemented with an inhibitor cocktail. The brains were collected at 3 or 24 h post-LPS treatment. Total RNA was extracted from MACS-sorted CD11b-enriched cell populations, whose purity, estimated by cytometry, ranged from 85.7 to 98.7% (**Fig. 5A**). As measured in tissue homogenates, LPS administration led to a significant inflammatory peak in microglial cells at 3 h, as reflected by high levels of *TNFα, IL-1β, IL-6, COX2, NOS2*, and *MCP1* transcripts (**Fig. 5B-G**). The calculated pro-inflammatory index peaked at 43.2-fold ± 6.4 of the control level (**Fig. 5H**).

**Figure 5.**
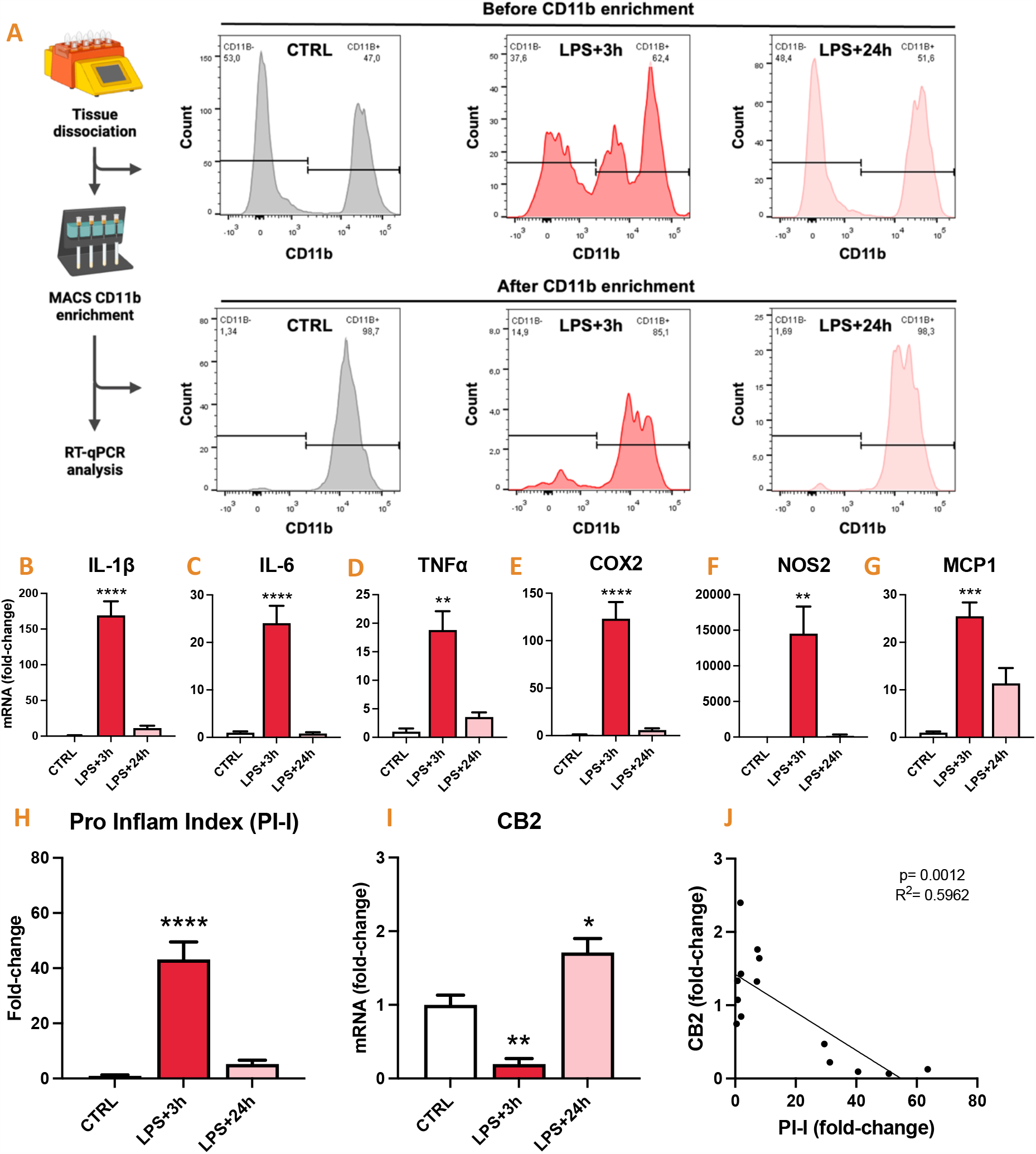
CB2 mRNA levels is inversely correlated with inflammatory markers in microglia after LPS administration in C57Bl/6 mice. Brain tissue of adult C57Bl/6 mice was dissociated using Miltenyi’s ABDK protocol and microglia-enriched cell population was sorted as in Fig. 2. Pro-inflammatory genes and CB2 mRNA were quantified by RT-qPCR in CD11b-positive cells sorted from mouse brain (neocortex and hippocampus) 3h (n=5) and 24h (n=5) following LPS administration (5 mg/kg, IP) and compared with untreated (CTRL) mice (n=4). **A**. The purity of the MACS-sorted CD11b-positive cells was estimated by quantifying the protein expression of CD11b by flow cytometry on a fraction of pooled cell suspensions harvested before and after CD11b MACS magnetic sorting. **B-G**. Quantification of IL-1β, IL-6, TNFα, COX2, NOS2 and MCP1 transcript levels. **H**. Pro-inflammatory index (PI-I) was calculated from transcript levels of pro-inflammatory genes (A-F). CTRL vs. LPS+3h: p<0.0001. **I**. Quantification of CB2 mRNA. CTRL vs. LPS+3h: p=0.0061; CTRL vs. LPS+24h: p=0.0132. Data are expressed as mean + SEM, and presented as relative to microglia from control mice. Normal data (IL-1β, IL-6, COX2, MCP1, PI-I and CB2) were analyzed with Tukey’s test following one-way ANOVA. Non-normal data (TNFα and NOS2) were analyzed with Dunn’s test following Kruskal-Wallis test. Asterisks represent LPS-treated groups vs CTRL: *, p<0.05; **, p<0.01; ***, p<0.001, ****, p<0.0001. **J**. Relationship between CB2-mRNA levels and PI-I (simple linear regression, p=0.0012, R2=0.5962, y=-0.02596x+1.423, n=14).

Three hours after LPS administration, CB2-mRNA levels in microglial cells transiently decreased to approximately five-fold that in controls, subsequently rebounding 24 h later to 171 ± 19% above that of controls once inflammation had resolved (**Fig. 5I**).

The CB2 transcript level was inversely correlated with the pro-inflammatory index after LPS administration (**Fig. 5J**). Conversely, GPR55 and GPR18 transcript levels did not significantly correlate with the pro-inflammatory index (**Fig. S6**).

### 3.6. CB2 mRNA levels in microglia BV2 cell line following LPS or IFNγ stimulation

Based on the quantification of pro-inflammatory gene transcripts (**Fig. 6A-F**) and calculation of the pro-inflammatory index (**Fig. 6G**), LPS treatment of murine BV2 microglial cells led to an inflammatory response that peaked between 2 and 4 h. Simultaneously, CB2-mRNA levels transiently decreased, reaching 12-fold lower values than those measured in untreated cells 2 h after LPS treatment (**Fig. 6H**). Considering the levels measured in brain tissue and sorted microglia, the CB2-mRNA level inversely correlated with the pro-inflammatory index (**Fig. 6I**). GPR55-mRNA levels were also inversely correlated with the pro-inflammatory index, whereas LPS-induced inflammation did not impact GPR18 expression (**Fig. S7**).

**Figure 6.**
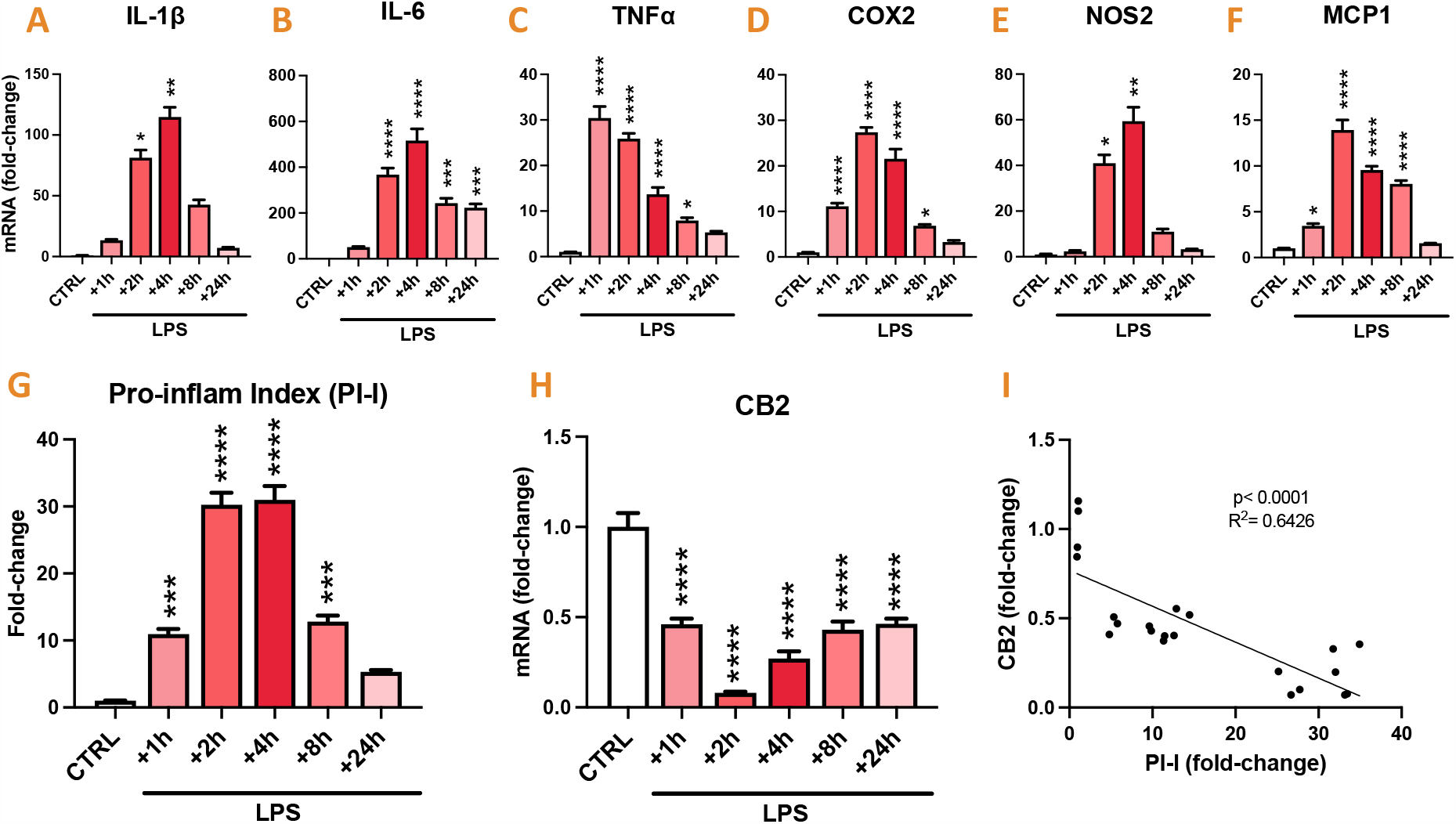
CB2 mRNA level is inversely correlated with that of inflammatory markers in BV2 cells after LPS treatment. Inflammatory genes and CB2 mRNA were quantified by calibrated RT-qPCR in cultured BV2 murine cells following LPS 1h, 2h, 4h, 8h or 24h application (100 ng/mL) and compared with levels quantified in untreated cells (CTRL, LPS+1h, +2h, +4h: n=4/condition; LPS+8h, +24h: n=3/condition). **A-F**. Quantification of IL-1β, IL-6, TNFα, COX2, NOS2 and MCP1 transcript levels. **G**. Pro-inflammatory index (PI-I) was calculated from transcript levels of pro-inflammatory genes (A-F). CTRL vs. LPS+1h: p=0.0005; vs. +2h: p<0.0001; vs. +4h, p<0.0001; vs. +8h: p=0.0002. **H**. Quantification of CB2-mRNA. CTRL vs. LPS+1h: p<0.0001; vs. +2h: p<0.0001; vs. +4h, p<0.0001; vs. +8h: p<0.0001; vs. +24h: p<0.0001. Data are expressed as the mean ± SEM, and presented as relative to untreated BV2 cells. Normal data (TNFα, IL-6, COX2, MCP1, PI-I and CB2) were analyzed with Tukey’s test for multiple comparisons following one-way ANOVA. Non-normal data (IL-1β and NOS2) were analyzed with Dunn’s test for multiple comparisons following Kruskal-Wallis test. Asterisks represent control (CTRL) vs. respective LPS-treated group. *, p<0.05; **, p<0.01; ***, p<0.001, ****, p<0.0001. **I**. Relationship between CB2-mRNA levels and PI-I (simple linear regression, p<0.0001, R2=0.6426, y=-0.02012x+0.7688, n=22).

Previous studies have shown that IFNγ-induced BV2 cell stimulation resulted in the reverse regulation of CB2 expression when compared with LPS induction ^12^. Komorowska-Müller et al. hypothesized that this difference in regulation could be attributed to a markedly low level of inflammation induced by IFNγ when compared with that induced by LPS and to the distinct nature of the two stimuli, i.e., IFNγ is a cytokine released during sterile inflammation, whereas LPS is a bacterial toxin ^7^. Treatment of BV2 cells with IFNγ for 3 or 8 h led to an increase in pro-inflammatory gene transcripts (**Fig. 7A-F**), as well as the pro-inflammatory index (**Fig. 7G**), which reached their highest values at 8 h. Simultaneously, CB2-mRNA levels did not decrease, but rather increased by up to 2-fold when compared with those of untreated cells (**Fig. 7H**) and were positively correlated with the pro-inflammatory index (**Fig. 7I**).

**Figure 7.**
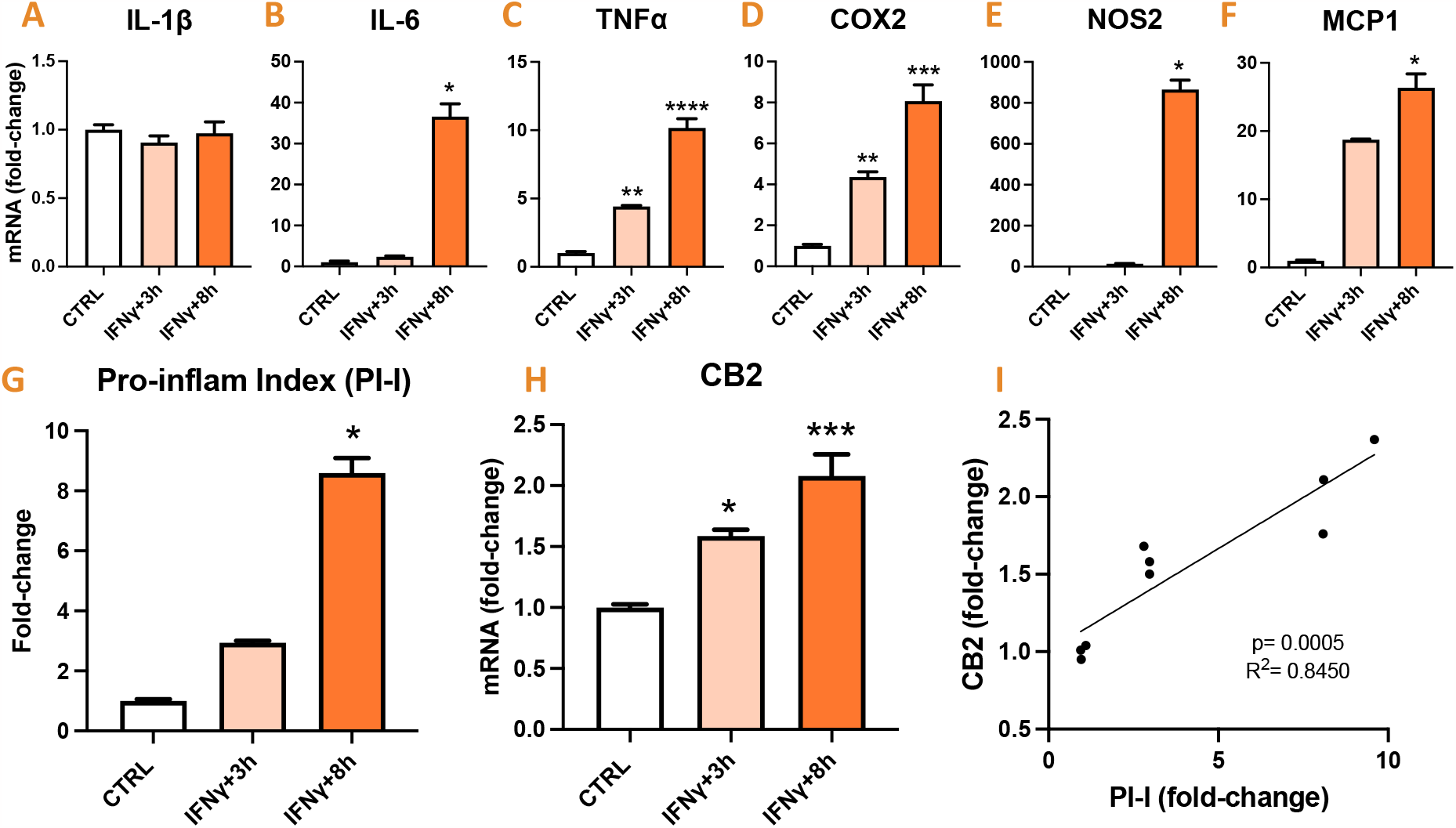
Level of CB2 mRNA positively correlated with that of inflammatory markers in BV2 cells after IFNγ treatment. Inflammatory genes and CB2 mRNA were quantified by calibrated RT-qPCR in cultured BV2 murine cells following IFNγ 3h or 8h application (100 ng/mL) and compared with levels quantified in untreated cells (n=3/condition). **A-F**. Quantification of IL-1β, IL-6, TNFα, COX2, NOS2 and MCP1 transcript levels. **G**. Pro-inflammatory index (PI-I) was calculated from transcript levels of pro-inflammatory genes (A-F). CTRL vs. IFNγ+8h: p=0.0211. **H**. Quantification of CB2 mRNA. CTRL vs. IFNγ+3h: p=0.0196; CTRL vs. IFNγ+8h: p=0.0010. Data are expressed as the mean ± SEM, and presented as relative to untreated BV2 cells. Normal data (IL-1β, TNFα, COX2 and CB2) were analyzed with Tukey’s post-hoc test for multiple comparisons following one-way ANOVA. Non-normal data (IL-6, NOS2, MCP1 and PI-I) were analyzed with Dunn’s post-hoc test for multiple comparisons following Kruskal-Wallis test. Asterisks represent control (CTRL) vs respective IFNγ-treated group. *, p<0.05; **, p<0.01; ***, p<0.001, ****, p<0.0001. **I**. Relationship between of CB2 transcript levels and pro-inflammatory index (simple linear regression, p=0.0005, R2=0.8450, y=0.1321x+1.005, n=9).

Interestingly, the stability of IL-1β-mRNA levels after IFNγ treatment in BV2 cells (**Fig. 7A**) was the main observed difference with LPS treatment. To determine whether the dramatic decrease in CB2-mRNA level, observed only under LPS treatment, could be attributed to the robust activation of IL-1R by IL-1β, BV2 cells were incubated with IL-1RA (1-1000 ng/mL), the natural IL-1R antagonist, 30-min prior to and during LPS. IL-1RA did not prevent the LPS-induced reduction in CB2 mRNA levels (**Fig. S8**).

## 4. DISCUSSION

### Main results

In the current study, we demonstrated that the CB2 receptor transcriptional level was consistently low, with uniform expression across various brain regions examined, including the olfactory bulb, neocortex, hippocampus, hypothalamus, cerebellum, and brainstem. Notably, there were no significant differences in CB2 expression between the three mouse strains (C57bl/6, Balb/c, and Swiss). Microglia were detected as the primary cell type expressing CB2 in the brain, with neurons displaying low levels of CB2 expression. In addition, we observed an inverse relationship between CB2 expression and LPS-induced inflammation over time, as evidenced by a reduction in CB2 transcript levels coinciding with the peak of inflammation, observed both at the tissue level in the hippocampus and neocortex and at the cellular level in sorted microglial populations and the BV2 microglial cell line. Intriguingly, we observed the opposite relationship after stimulation of BV2 cells with IFNγ, which activates distinct intracellular pathways from LPS.

### Methodological consideration

One objective of the current study was to identify the cell types that contribute to CB2 expression in the brain tissue using cell-sorting protocols. Obtaining a cell suspension from the brain tissue and enriching cell populations by magnetic or fluorescent sorting represents an assault on brain cells that can alter their phenotypic state. Microglia are effective sentinels that continuously scan the microenvironment in their basal state, entering an extremely rapid activation process whenever cerebral homeostasis is compromised ^18,19^. *Ex vivo* microglial activation can introduce confounders that can distort measurements of the transcriptomic profile in the basal state or mask endogenously induced activation, similar to pathological conditions ^17^. Previous studies have shown that supplementing the buffer with transcription and translation inhibitors during the enzymatic and mechanical dissociation of brain tissue limits the *ex vivo* activation of microglial cells ^10^. Herein, we showed that using the same inhibitors from the intracardiac perfusion stage through to final cell collection could limit the induction of IL-1β and TNFα pro-inflammatory markers in microglial populations enriched from brain tissue of healthy adult mice. As sentinel cells of the CNS, microglial cells are most likely to be sensitive to *ex vivo* activation during cell dissociation and sorting. However, it should be noted that uncontrolled gene activation may also occur in other cell types during these steps. We only examined the effect of inhibitors on microglial cells and not on other cell populations. We assumed that inhibiting the non-physiological activation of microglia may also extend to other brain cell types, given that targets of transcriptional and translational inhibitors are conserved in all eukaryotic cell types ^20,21^ and are, therefore, not cell-specific.

### CB2 basal expression in the CNS

Under physiological conditions, we did not detect inter-strain (C57bl/6, Balb/c, and Swiss) or any spatial regulation of *Cnr2* gene expression in the examined brain regions: the olfactory bulb, neocortex, hippocampus, hypothalamus, cerebellum, and brainstem. CB2 transcript levels were relatively low when compared with those of the other main cannabinoid receptors, i.e., CB1, GPR18, and GPR55. Confirming the presence of CB2 in the physiological state is consistent with quantifiable behavioral outcomes in healthy mice, such as memory ^22,23^, mood ^24^ or pain sensitivity ^25,26^ measured after administration of CB2-specific ligands ^5^. The homogeneity of CB2 expression contrasted with the heterogeneity between the investigated regions, both in terms of structural organization and function. This finding suggests that CB2 is predominantly expressed by cells with a supporting role in the CNS rather than by those with a highly specialized role. Although subtle differences in density and phenotype have been identified in certain brain regions, microglial cells are present throughout the nervous system and play crucial homeostatic roles ^27–29^. Furthermore, CB2 is mostly expressed in the periphery by leukocytes, including myeloid monocytes and macrophages ^1,30,31^, suggesting that microglia are one of the main cells expressing CB2 in the CNS.

In the present study, we used MACS to investigate the CB2 expression in enriched populations sorted from adult mouse brains. The MACS technique enabled us to obtain yields required for downstream transcript analysis in a short period. We showed that CB2 was primarily expressed in the brain by CD11b-positive microglial cells and, to a lesser extent, by neurons under basal conditions. However, MACS enrichment based on the use of a single-cell marker is insufficient to distinguish the subpopulations present in a heterogeneous population. Upon confirming that CB2 expression in the neocortex and hippocampus of 10-day-old mice was identical to that in adult mice, we analyzed single nuclei from the cortex of 10-day-old mice to gain a better understanding of the distribution of cells expressing CB2-mRNA in the neocortex. Under basal conditions, less than 1% of cortical cells express CB2. Furthermore, this analysis revealed that microglia constituted the highest proportion of CB2-positive cells, consistent with the MACS results obtained in adult mice.

These results provide new insights into the cells involved in physiological and behavioral outcomes observed following the pharmacological activation of CB2 in healthy mice. Neuronal CB2 may directly modulate behavioral outcomes ^4,5^. Microglia have been shown to play an indirect role in neuronal activity and regulation of behavioral outcomes ^32^. Microglial CB2 may thus, in part, be responsible for certain outcomes reported in the physiological state.

### CB2 expression in activated microglia

Numerous studies have reported that CB2 is a target molecule in microglial cells and monocytes in different models of pathological conditions for efficiently resolving neuroinflammation ^5,7,33,34^. In addition, while CB2 transcript levels have been found to increase under numerous pathological conditions at the brain tissue level ^12,35–38^, it has been speculated that this induction mainly occurs in microglial cells ^12,37,39^. We elicited robust evidence that CB2 expression is inversely regulated by the LPS-induced inflammatory state, both at the tissue level in microglial cells sorted from the brain tissue and in the BV2 microglial cell line, and is induced during the resolution phase of inflammation rather than during the inflammatory peak. These results are consistent with the few studies that have demonstrated that CB2 transcript levels are downregulated in microglia cell lines stimulated with LPS ^11,12^ or LPS+IFNγ ^13,40^. However, they contradicted the general idea that CB2 is induced in the brain in many pathological models of neuroinflammation ^7,9^. One potential explanation is the distinct nature of the inflammatory stimulus, which may have different effects on the regulation of CB2 expression by engaging various intracellular pathways. The stimulation of BV2 cells with IFNγ was found to yield an opposite regulation of CB2 expression when compared to the effect induced by LPS. However, the inflammatory responses elicited by LPS and IFNγ were not assessed, rendering it challenging to ascertain whether this discrepancy stemmed from variations in the intensity of inflammation or the characteristics of the inflammatory response ^12^. LPS mimics non-sterile inflammation by activating the Toll-like receptor 4 (TLR4) and numerous subsequent intracellular pathways, including NF-κB ^41^, while IFNγ simulates sterile inflammation and binds to the interferon-gamma receptor (IFNGR) protein complex, which activates intracellular JAK/STAT pathways ^42^. In the current study, we showed that CB2 transcript levels exhibited an inverse regulation in BV2 cells when stimulated with a similar dose of LPS and IFNγ. We demonstrated that BV2 cells exhibited distinct responses to LPS and IFNγ, with differences in terms of the temporal pattern, magnitude, and cytokine profile. The IFNγ-induced inflammatory response had a slow onset, low magnitude, and did not result in the upregulation of IL-1β transcript levels. To investigate whether the substantial decrease in CB2 mRNA levels, observed exclusively with LPS treatment, was a consequence of potent IL-1R activation by IL-1β, we stimulated BV2 cells with LPS in the presence of the natural IL-1R antagonist IL-1RA. Interestingly, we observed no difference in the LPS-induced decrease in CB2 mRNA levels, suggesting that IL-1β does not drive LPS-induced CB2 downregulation. Further investigations are necessary to clarify these mechanisms.

Notably, the upregulation of CB2 transcripts observed in the brain of several experimental models linked to neuroinflammatory processes can also be attributed to the infiltration of circulating leukocytes into the brain parenchyma, which are known to exhibit robust CB2 expression ^1^. into the brain parenchyma. Substantial upregulation of CB2 at the transcriptional level has been documented in the brain tissues of rodent models of stroke ^38^, traumatic brain injury ^35^ and Parkinson’s disease ^36,37^, in which strong leukocyte infiltration has been reported ^35,43–46^.

## Conclusion

These results represent a substantial advancement in our understanding of CB2 expression and its role in the CNS under both physiological and pathological conditions. Our findings highlight that CB2 expression is differentially regulated in distinct inflammatory environments. Therefore, it is mandatory to measure CB2 expression in each experimental model before considering pharmacological interventions to identify precise target cells and optimal therapeutic windows.

## Supporting information

Method details

## 5. ACKNOWLEDGMENTS

We acknowledge the contribution of SFR Santé Lyon-Est (UAR3453 CNRS, US7 Inserm, UCBL) CyLE cytometry platform facilities, especially Thibault Andrieu and Priscillia Battiston-Montagne for their valuable help. We are grateful for animal care provided by the zootechnicians of the CRNL’s animal facility. Nadia Gasmi was granted a PhD fellowship from the Fondation pour la Recherche Médicale. Wanda Grabon was granted a PhD fellowship from France Alzheimer.

## 6. AUTHOR’S CONTRIBUTION

LB and WG conceived and designed the study. WG, AR, NG, BG, SR, AB and JB participated in data collection and analysis. GM and CD performed RNAseq single nucleus experiments and analysis. WG, LB and GM interpreted the data. WG drafted the manuscript. LB provided critical revisions. All authors read and approved the final manuscript.

## HIGHLIGHTS

- CB2 receptor mRNA expression was low and uniform across various brain regions examined.
- The use of transcription and translation inhibitors during brain dissociation and cell sorting efficiently prevented *ex vivo* microglial activation.
- CB2 mRNA was detected in a few cells in the physiological state and was mainly detected in microglia and some neurons.
- CB2 expression is downregulated in microglia during the LPS-induced inflammatory peak and upregulated during the resolution of inflammation.
- LPS and IFNγ stimulation distinctly regulate CB2 expression in microglia.

