## Supplementary material for "Cannabinoid receptor type 2 expression in mouse brain: from mapping to regulation in microglia under inflammatory conditions": Method details

### Methods details

#### Animals

Adult male mice (Balb/c, C57Bl/6 and Swiss, 8-weeks-old, Envigo, France) were used in this study. Animals were maintained under a constant light/dark cycle (12:12 h, lights on at 07:00 a.m.) with *ad libitum* access to food and water. Room temperature was kept at 23±1°C. The experimental procedures were conducted in accordance with the European Community guidelines for care in animal research and approved by the CELYNE local Ethics Research Committee (protocol #24302). Every effort was made to minimize animal suffering.

#### BV2 Cell culture

The immortalized murine BV2 cell line (BV2 cells) was kindly provided by Dr. Nadia Soussi (NeuroDiderot, Paris University) and cultured in high-glucose Dulbecco's Modified Eagle's medium (DMEM GlutaMAX-I, Gibco Life technologies #10566-016), supplemented with 10% heat-inactivated fetal bovine serum (FBS, Gibco Life technologies #10270-098), and 1% antibiotic-antimycotic (Gibco Life technologies #15240-096). Cells were cultured at a density of 1 x $10^5$/$cm^2$ in 6-wells plates in a humidified incubator of 5% $CO_2$ at 37°C. At P9, cells were treated with LPS (from *Escherichia coli* O55:B5, 100 ng/mL, Sigma #L6529) for 1h, 2h, 4h, 8h or 24h, or with IFNγ (100 ng/mL, Gibco #PMC4031) for 3h or 8h, and harvested for further RT-qPCR analysis. For the study aimed at tackling the role of IL-1R signaling in CB2 mRNA decrease following LPS (100 ng/mL), BV2 cells were pre-treated for 30 min with recombinant mouse IL1-Ra protein (1-1000 mg/mL, abcam #ab283475), and then co-incubated with LPS + IL1-Ra for 2h.

#### RNA extraction

Tissues (Experiments 1 & 3) were crushed in 250 µL of molecular biology grade water (Eurobio) using Tissue-Lyser II (Qiagen) according to the manufacturer's instructions. Total RNAs from brain structures and from BV2 cells (Experiment 4) were extracted using Tri-Reagent LS (Molecular Reasearch Center, #TS120) or Tri-Reagent (Molecular Reasearch Center, #TR118), respectively. Genomic DNA was removed using Turbo-DNA-free (Ambion, #1907) and total RNAs were purified with RNeasy mini kit (Qiagen, #74104). For sorted cells (Experiment 2 & 4), total RNAs were extracted and purified using the RNeasy Plus Micro Kit (Qiagen, #74034) according to manufacturer’s instruction. RNA concentration was determined for each sample on the BioDrop® µLite.

**Reverse transcription and real-time quantitative PCR**

Total tissue RNAs were reverse transcribed to complementary DNA (cDNA) using both oligo dT and random primers with PrimeScript RT Reagent Kit (Takara, #RR037A) according to manufacturer's instructions, in a total volume of 10 µL. In RT reaction, 300,000 copies of a synthetic external non-homologous poly(A) standard messenger RNA (SmRNA; A. Morales and L. Bezin, patent WO2004.092414) were added to normalize the RT step, as previously described [16]. cDNA was diluted 1:13 with nuclease free Eurobio water and stored at -20°C until the further use. Each cDNA of interest was amplified on 5 µL of the diluted RT reaction by real-time PCR, using the Rotor-Gene Q thermocycler (Qiagen), the SYBR Green PCR kit (Qiagen, #208052) and oligonucleotide primers (Eurogentec) specific to the targeted cDNA. The sequences of the specific forward and reverse primer pairs were constructed using Primer-BLAST (NCBI). Primers used are listed in **Table S1**. cDNA copy number detected was determined using a calibration curve, and results were expressed as cDNA copy number / µg tot RNA.

Pro-inflammatory index was calculated using a specific set of pro-inflammatory genes: IL-1β, IL-6 and TNFα. For each sample, the number of copies of each transcript has been expressed in percent of the averaged number of copies measured in the whole considered group of samples. Once each transcript is expressed in percent, an index was calculated by adding the percent of each transcript involved in the composition of the index and expressed in arbitrary units (A.U.).

**Brain dissociation and magnetic cell sorting (MACS studies)**

To prevent any artifactual *ex vivo* gene expression changes during brain dissociation and cell sorting procedures, all buffers and solutions used during the process (from animal perfusion to sorted cell flash freezing) were supplemented with a cocktail composed of Actinomycin D (3µM, Tocris #1229/10), Anisomycin (100 µM, Tocris #1290/50) and Triptolide (10 µM, Tocris #3253/10) i.e. transcription and translation inhibitors [10]. All steps were performed on ice or using pre-chilled refrigerated centrifuge set to 4°C with all buffers/solutions pre-chilled before addition to samples to further limit cell activation. Buffer 1 (B1) was composed of DPBS 1X (Thermofisher #14040-117) and inhibitor cocktail. Buffer 2 (B2) was composed of DPBS 1X, BSA 0,5% (Sigma #A2153) and inhibitor cocktail. The general workflow of brain dissociation and magnetic cell sorting is illustrated in figure 1.

**Brain dissociation** – Mice were anesthetized and perfused intracardially with ice-cold B1. Neocortices and hippocampi were quickly dissected and placed in B1 until all perfusions were completed. Tissues were then cut in smaller pieces with a scalpel and processed for dissociation using Miltenyi's Adult Brain Dissociation Kit (#130-107-677) according to manufacturer's instruction. Inhibitor cocktail was added in each reagent. Briefly, samples were added to gentleMACS C Tubes (Miltenyi #130-093-237) with the enzyme mixes and placed in gentleMACS OctoDissociatior with heaters (Miltenyi #130-096-

427), running program 37C_ABDK_01. Once program finished, samples were briefly spun before being filtered through 70 µm cell strainer (Thermofisher #11597522). Samples were washed with B1 and spun to pellet cells. To clear cell solution, cell pellets were resuspended and overlaid with appropriate volume of Miltenyi Debris Removal Solution according to manufacturer's protocol. Debris were removed from top layer and solution was diluted with B1 and spun to pellet cells. Cells collected were resuspended in B1 and counted manually (with trypan blue) before magnetic sorting.

**Magnetic Cell-sorting –** To enhance cell yields, distinct mice were used for the isolation of neurons (n=3) and for the isolation of the other cell types (microglia, endothelial cells and astrocytes, n=3). Neurons were enriched using Adult Neuron Isolation Kit (Miltenyi #130-126-602) according to manufacturer's instructions. Briefly, brain cells from the micro dissected hippocampi and neocortices were incubated with Adult Non-Neuronal Cell Biotin-Antibody Cocktail, washed with B2, incubated with Anti-Biotin MicroBeads, volume was adjusted with B2 and solutions were applied onto LS columns (Miltenyi #130-042-401) in the magnetic field of QuadroMACS Separator (Miltenyi #130-090-976). To improve yields and purity of cell suspensions, unlabeled cells were applied a second time on new LS columns. Unlabeled cells (i.e., neuronal cells) collected were then counted manually, spun and dry cells pellets were flash frozen and stored at -80°C for further analysis. In distinct mice, endothelial cells, microglia and astrocytes were magnetically sorted successively. Cells were first incubated with mouse FcR Blocking Reagent (Miltenyi #130-092-575). Endothelial cells were first enriched using anti-Ly-6C biotin antibody (Miltenyi #130-111-914). Following antibody incubation, cells were washed with B2, spun, resuspended in B2 and incubated with Anti-Biotin MicroBeads (Miltenyi #130-090-485). Cells were then washed, spun and resuspended in 500µL of B2. Cell suspensions were applied onto MS columns (Miltenyi #130-042-201) in the magnetic field of OctoMACS Separator (Miltenyi #130-042-109). To improve purity of cell suspensions, the positive fractions were applied a second time on new MS columns. Labeled cells (i.e., endothelial cells) collected were then counted manually, spun and dry cells pellets were flash frozen and stored at -80°C for further use. Unlabeled cells were resuspended in B2 to proceed to Microglial enrichment using CD11b MicroBeads (Miltenyi #130-093-634). Following incubation, cells were washed, spun and resuspended in 500µL of B2. Magnetic separation was performed as for endothelial cells enrichment twice. Labeled cells (i.e., microglial cells) collected were then counted manually, spun and dry cells pellets were flash frozen and stored at -80°C. Unlabeled cells were resuspended in B2 to proceed to astrocytes enrichment using ACSA-2 MicroBeads (Miltenyi #130-097-679). Following incubation, cells were magnetically sorted as for endothelial and microglial cells, twice. Labeled cells (i.e., astrocytes) collected were then counted manually, spun and dry cells pellets were flash frozen and stored at -80°C.

**Flow Cytometry**

To control purity of enriched cell populations, a fraction of cell suspensions was collected before and after each sorting. The following antibodies were added to cell suspensions at 1:50 concentration in B2 buffer: ACSA2-APC (Miltenyi #130-116-245) CD11b-PE Vio770 (Miltenyi #130-113-808), CD31-PE (Miltenyi #130-111-540), Ly6C-APC Vio770 (Miltenyi #130-111-919) and O4-VioBright515 (Miltenyi #130-120-102). Samples were incubated with staining antibodies for 15 min at 4°C; washed in B2 and then spun down for 10 min at 300g, before being resuspended in 100µL of B2. DAPI (2.5 µg/mL, Sigma #D9542) was added as a viability marker right before flow cytometry analysis. Cells were analyzed with the BD FACS Canto II Flow Cytometer (Becton-Dickinson) and data files with FlowJo software V10.7.2 (Becton-Dickinson). The same gating strategy has been used for all data files, Side scatter-A (SSC-A) and Forward scatter-A (FSC-A) were used for cells morphology gating, FSC-W and FSC-H as well as SSC-W and SSC-H were used to gate singlets and DAPI- cells for alive cells. Then each specific markers were used to detect population of interest, $CD11b^+$ for microglial cells, $O4^+$ for oligodendrocytes, $ACSA2^+$ for astrocytes, $Ly6c^+$ / $CD31^+$ for endothelial cells and all these negative markers for neuronal cells.

**Statistical Analysis**

Statistical analyses were performed using Prim 10.0 software (GraphPad, USA). Results are presented as mean ± SEM (standard error of the mean). Differences with a p-value<0.05 ($p<0.05$) were considered to be statistically significant. The Shapiro–Wilk test and quantile–quantile plot were used to assess normal distribution of the data. For normal data, the statistical significance was assessed by t-test or one-way ANOVA analysis, followed with Tukey's post-hoc test for multiple comparisons. For non-normal data, the statistical significance was assessed by Kruskal-Wallis test, followed with Dunn's post-hoc test for multiple comparisons. Simple linear regression was used to assess the association between the level of expression of cannabinoid receptors genes and pro-inflammatory index.
