## Supplementary material for "Cannabinoid receptor type 2 expression in mouse brain: from mapping to regulation in microglia under inflammatory conditions": Figure S1

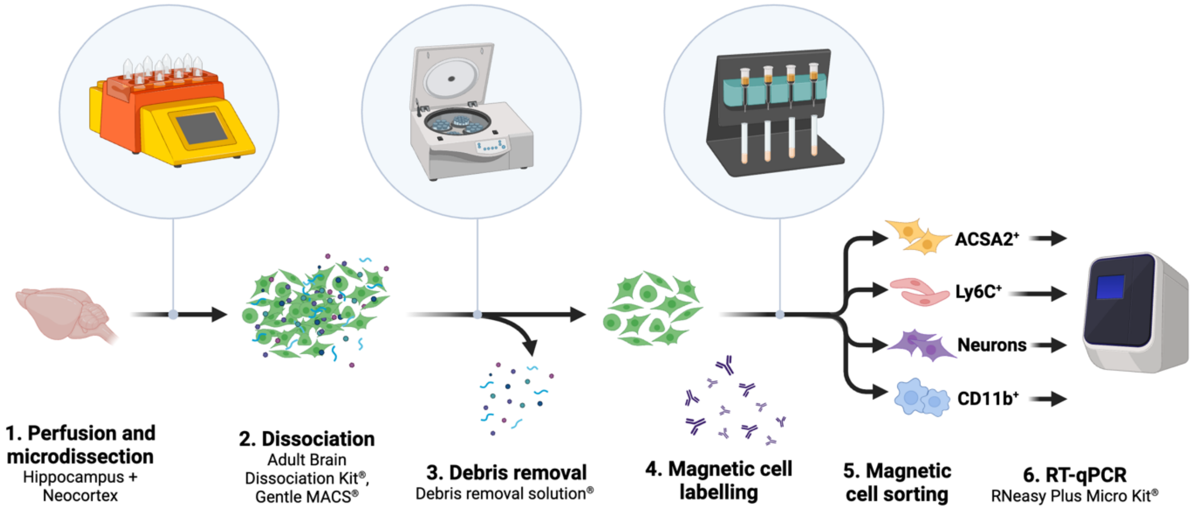


**Figure S1 – Brain dissociation and magnetic cell sorting workflow.**
