## Supplementary material for "Cannabinoid receptor type 2 expression in mouse brain: from mapping to regulation in microglia under inflammatory conditions": Figure S2

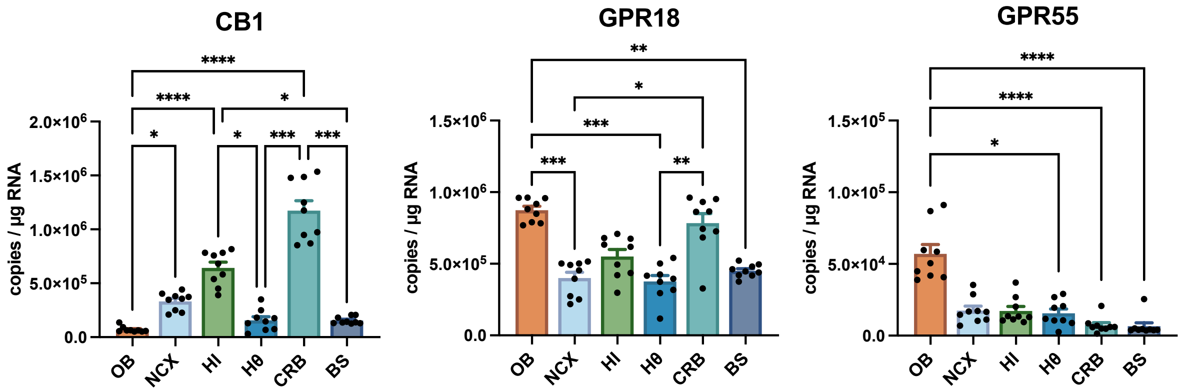


**Figure S2 – Regional distribution of CB1, GPR18 and GPR55 cannabinoid receptors mRNA in the healthy mouse brain.** CB1 (**A**), GPR18 (**B**) and GPR55 (**C**) mRNA was quantified using calibrated RT-qPCR in six microdissected regions from adult mouse brain: the olfactory bulb (OB), the neocortex (NCX), the hippocampus (HI), the hypothalamus (Hθ), the cerebellum (CRB) and the brainstem (BS), n=9 mice. Transcript levels are expressed as copies of cDNA per microgram of total RNA. Values are presented as means + SEM. Data were analyzed with Dunn's post-hoc test for multiple comparisons following Kruskal-Wallis test. *, p<0.05; **, p<0.01; ***, p<0.001, ****, p<0.0001. Not significantly statistical differences are not graphically represented.
