## Supplementary material for "Cannabinoid receptor type 2 expression in mouse brain: from mapping to regulation in microglia under inflammatory conditions": Figure S3

**A. ACSA2 enrichment (astrocytes/oligodendrocytes)**

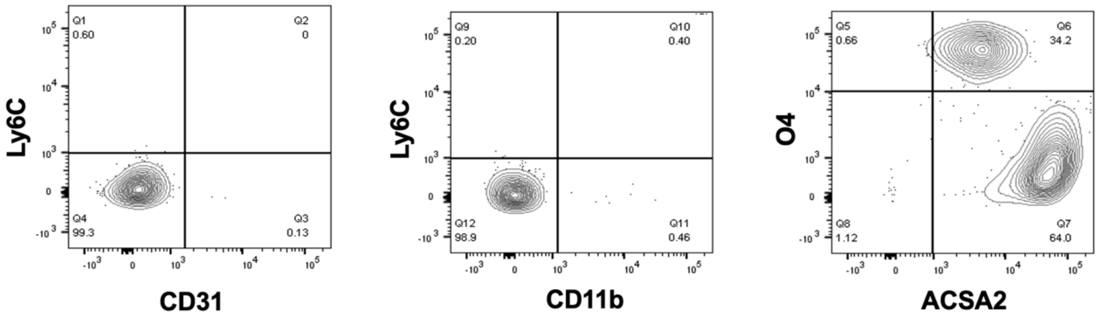


**B. CD11b enrichment (microglia)**

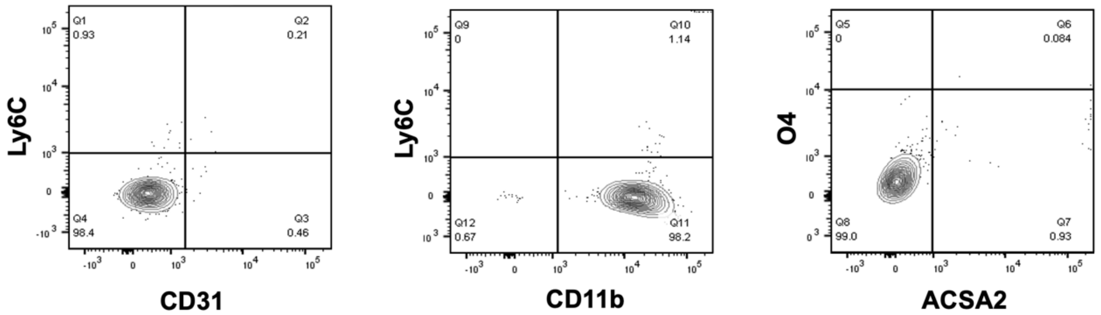


**C. Ly6C enrichment (endothelial cells)**

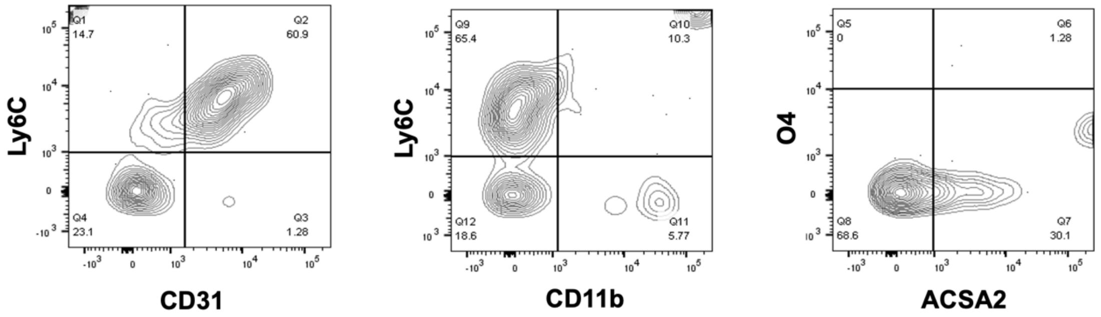


**D. Neuron isolation**

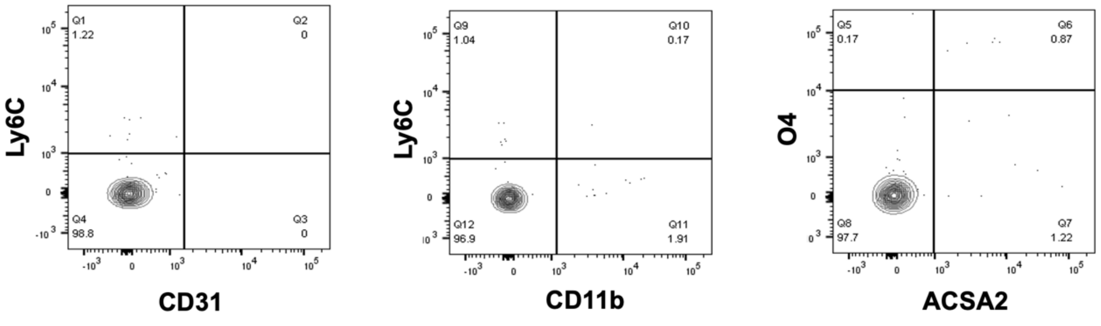


**Figure S3** – **Purity of MACS enriched cell populations from healthy mouse brain.** The purity of the MACS-sorted cell populations was estimated by quantifying the protein expression of specific cell markers (CD11b, Ly6C, O4, ACSA2, and CD31) by flow cytometry on a fraction of pooled cell suspensions harvested after each MACS magnetic sorting. **A**. Cells sorted using anti-ACSA-2 antibody (Miltenyi # 130-097-679). Flow cytometry data revealed that both astrocytes (ACSA2$^{+}$ O4$^{-}$) and oligodendrocytes (ACSA2$^{+}$ O4$^{+}$) have been enriched in this cell fraction. **B**. Cells sorted using anti-CD11b antibody (Miltenyi #130-093-634). **C**. Cells sorted using anti-Ly6C antibody (Miltenyi #130-111-914). **D**. Cells sorted using Adult Neuron Isolation Kit (Miltenyi #130-126-602).
