## Supplementary material for "Cannabinoid receptor type 2 expression in mouse brain: from mapping to regulation in microglia under inflammatory conditions": Figure S4

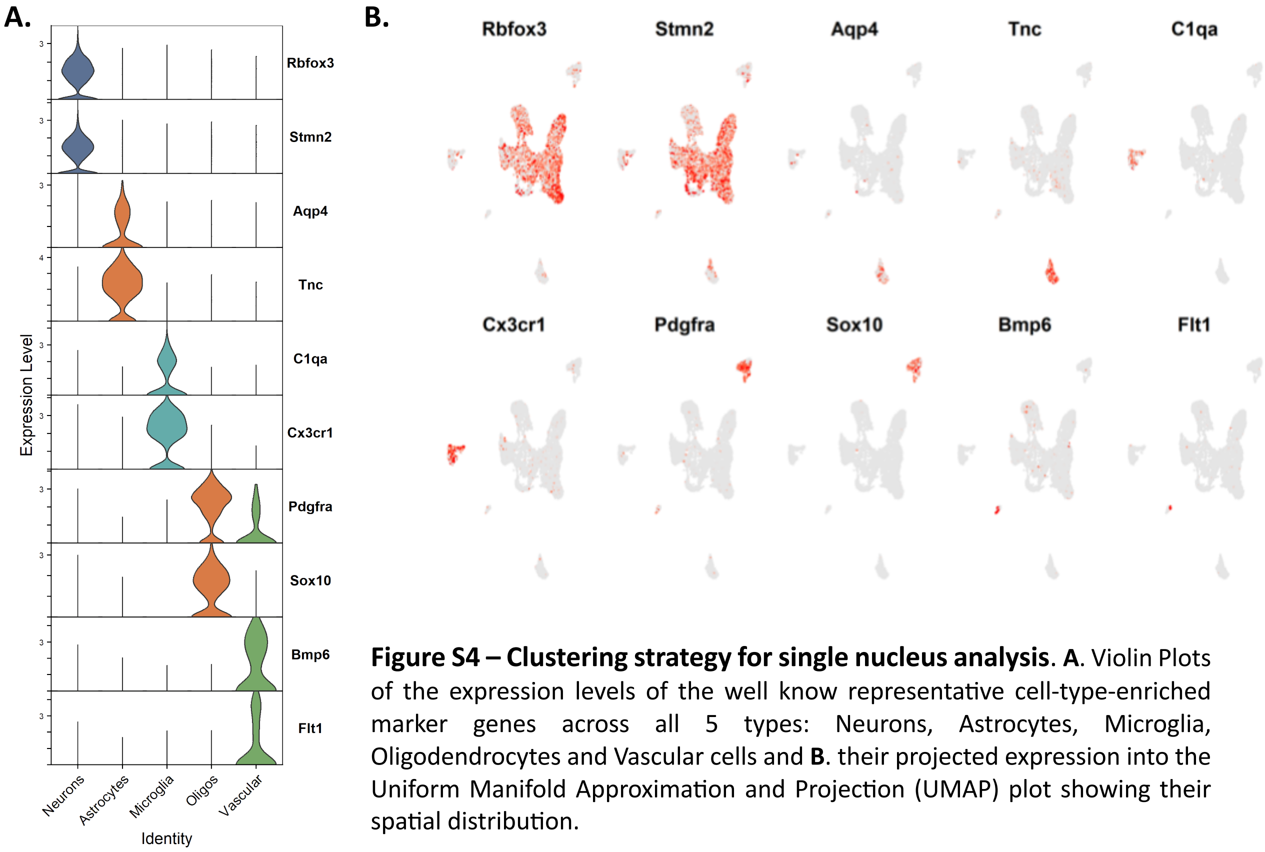


**Figure S4 – Clustering strategy for single nucleus analysis**. **A**. Violin Plots of the expression levels of the well know representative cell-type-enriched marker genes across all 5 types: Neurons, Astrocytes, Microglia, Oligodendrocytes and Vascular cells and **B**. their projected expression into the Uniform Manifold Approximation and Projection (UMAP) plot showing their spatial distribution.
