## Supplementary material for "Cannabinoid receptor type 2 expression in mouse brain: from mapping to regulation in microglia under inflammatory conditions": Figure S5

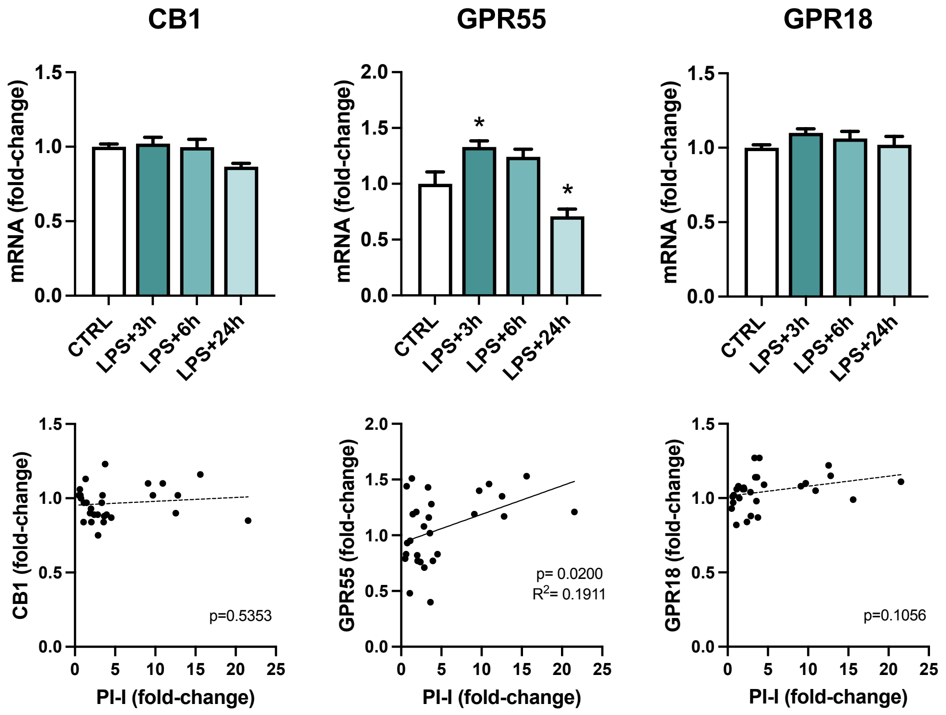


**Figure S5 – Expression of other cannabinoid receptors in the brain during LPS-induced inflammation.** **A.** CB1, GPR55 and GPR18 mRNA were quantified using calibrated RT-qPCR in the brain of adult mouse brain 3h (n=7), 6h (n=7) and 24h (n=8) following LPS administration (5 mg/kg, IP) and compared with levels quantified in untreated mice (n=6). Data were analyzed with Tukey's post-hoc test following one-way ANOVA. Asterisks represent control (CTRL) vs respective LPS-treated group. *, $p<0.05$; **, $p<0.01$; ***, $p<0.001$, ****, $p<0.0001$ **B**. Relationship between CB1, GPR55 and GPR18 transcript levels and pro-inflammatory index (simple linear regression, CB1: $p=0.5353$; GPR55: $p=0.0200$, $R^2=0.1911$, $y=0.02565x+0.91314$; GPR18: $p=0.1056$), n=28.
