## Supplementary material for "Cannabinoid receptor type 2 expression in mouse brain: from mapping to regulation in microglia under inflammatory conditions": Figure S6

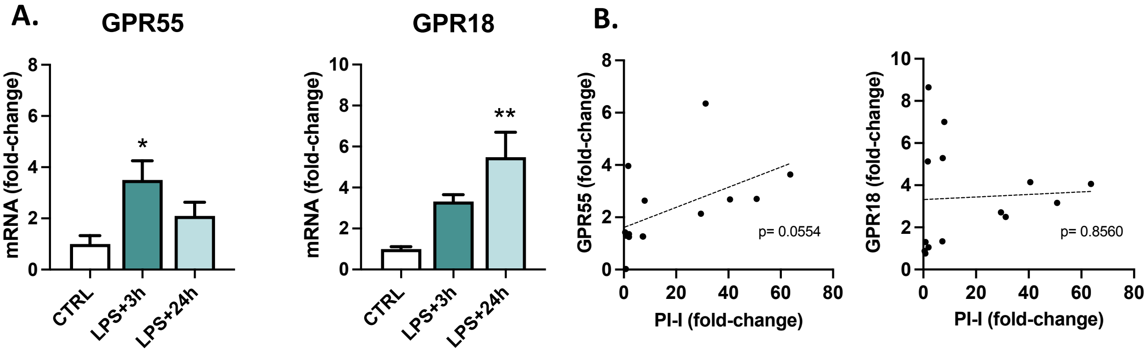


**Figure S6 – Expression of other cannabinoid receptors in microglia during LPS-induced inflammation.** **A.** GPR55 and GPR18 mRNA were quantified using calibrated RT-qPCR in CD11b-positive cells sorted from adult mouse brain 3h (n=5) and 24h (n=5) following LPS administration (5 mg/kg, IP) and compared with levels quantified in untreated mice (n=4). CB1 mRNA has not been quantified as CB1 is almost undetectable in microglial cells. Transcript levels are expressed as relative levels of control mice. Values are presented as means ± SEM. Normal data (GPR18) were analyzed with Tukey's post-hoc test for multiple comparisons following one-way ANOVA. Non-normal data (GPR55) were analyzed with Dunn's post-hoc test for multiple comparisons following Kruskal-Wallis test. Asterisks represent control (CTRL) vs respective LPS-treated group. *, p<0.05; **, p<0.01; ***, p<0.001, ****, p<0.0001 **B**. Relationship between GPR55 and GPR18 transcript levels and pro-inflammatory index (simple linear regression, GPR55: p=0.0554, GPR18: p=0.8560).
