## Supplementary material for "Cannabinoid receptor type 2 expression in mouse brain: from mapping to regulation in microglia under inflammatory conditions": Figure S7

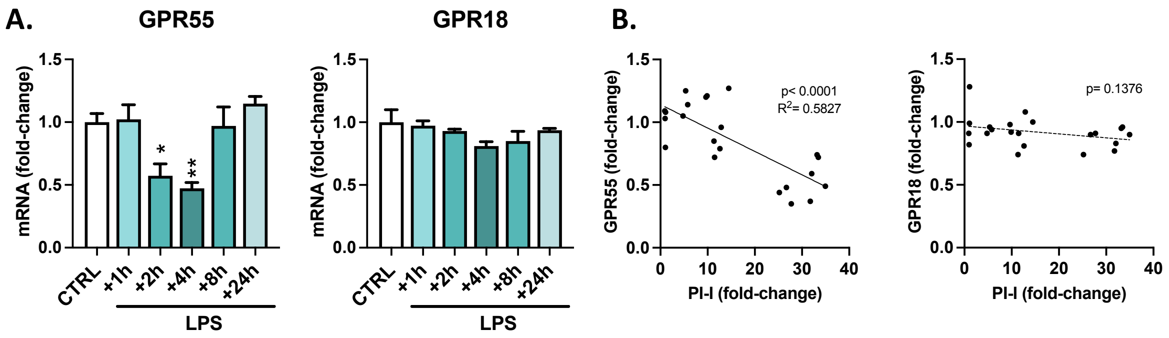


**Figure S7 – Expression of other cannabinoid receptors in BV2 cells during LPS-induced inflammation.** **A.** GPR55 and GPR18 mRNA were quantified using calibrated RT-qPCR in murine BV2 microglial cells 1h, 2h, 4h, 8h or 24h following LPS application (100 ng/mL) and compared with levels quantified in untreated cells (CTRL, LPS+1h, +2h, +4h: n=4/condition; LPS+8h, +24h: n=3/condition). CB1 mRNA has not been quantified as CB1 is almost undetectable in microglial cells. Transcript levels are expressed as relative levels of untreated cells. Values are presented as means ± SEM. Normal data (GPR55 and GPR18) were analyzed with Tukey's post-hoc test for multiple comparisons following one-way ANOVA. Asterisks represent control (CTRL) vs respective LPS-treated group. *, $p<0.05$; **, $p<0.01$; ***, $p<0.001$, ****, $p<0.0001$ **B**. Relationship between GPR55 and GPR18 transcript levels and pro-inflammatory index (simple linear regression, GPR55: $p<0.0001$, $R^2=0.5827$, $y=-0.01865x+1.141$, GPR18: $p=0.1376$), n=22.
