## Supplementary material for "Cannabinoid receptor type 2 expression in mouse brain: from mapping to regulation in microglia under inflammatory conditions": Figure S8

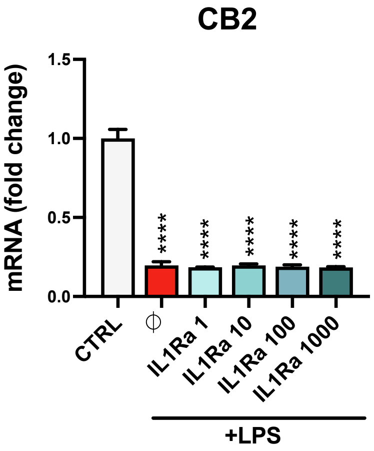


**Figure S8 – Blockade of IL-1R with IL1Ra treatment did not prevent LPS-induced CB2 mRNA decrease in BV2 cells.** CB2 mRNA was quantified by calibrated RT-qPCR in cultured BV2 murine cells following 2h LPS stimulation (100 ng/mL), with or without IL-1Ra (1-1000 ng/mL) pre-treatment and compared with levels quantified in untreated cells (n=3/condition). IL1Ra was applied 30min before LPS challenge. Data are presented as means + SEM, and were analyzed with Tukey's post-hoc test for multiple comparisons following one-way ANOVA. Asterisks represent control (CTRL) vs respective LPS-treated group. *, $p<0.05$; **, $p<0.01$; ***, $p<0.001$, ****, $p<0.0001$
