## Supplementary material for "Cannabinoid receptor type 2 expression in mouse brain: from mapping to regulation in microglia under inflammatory conditions": Table S1

**Table S1 – Sequences of primer pairs used for qPCR** (*Mus Musculus*)

| Target cDNA | Forward primer sequence | Reverse primer sequence | GenBank Reference |
|---|---|---|---|
| **CB1** | 5’ CCA GCA GGA GAC ACA ACC AA 3’ | 5’ CTG CGG GAG TGA AGG ATG AC 3’ | NM_001355021.2 |
| **CB2** | 5’ AAG GCC AGA TCT CCT CTC AC 3’ | 5’ TCA CTT CTG TCT CCC GGC AT 3’ | NM_009924.4 |
| **CD31** | 5’ GCA TCG GCA AAG TGG TCA AG 3’ | 5’ TGT TGC TGG GTC ATT GGA GG 3’ | NM_001032378 |
| **CD115** | 5’ AAT CCA CCT CCA CTG GCA TC 3’ | 5’ TTG AAT CCC ACT TCG GCG TT 3’ | NM_001037859.2 |
| **COX2** | 5’ AGC CCA TTG AAC CTG GAC TG 3’ | 5’ ACC CAA TCA GCG TTT CTC GT 3’ | NM_011198.4 |
| **GFAP** | 5’ GGC TCG TGT GGA TTT GGA GA 3’ | 5’ GCC ACT GCC TCG TAT TGA GT 3’ | NM_001131020.01 |
| **GLAST** | 5’ CTG GTA ACC CGG AAG AAC CC 3’ | 5’ GGG GAG CAC AAA TCT GGT GA 3’ | NM_148938 |
| **GPR18** | 5’ TAA TGG CTC ACA CCC AGA GG 3’ | 5’ AAA ACA TCC GAA AGG GCA GAC 3’ | NM_182806.2 |
| **GPR55** | 5’ GTG TGA AGC AGG TTG TGT GG 3’ | 5’ CTG TCC AAG TCA GAA GGC TGG 3’ | NM_001033290.2 |
| **Iba1** | 5’ GGC TGG AGG GGA TCA ACA AG 3’ | 5’ GAG TAG CTG AAC GTC TCC TCG 3’ | NM_001361501 |
| **IL-1β** | 5’ GCT GAA AGC TCT CCA CCT CA 3’ | 5’ CTT GGG ATC CAC ACT CTC CAG 3’ | NM_008361.3 |
| **IL-6** | 5’ TCC GGA GAG GAG ACT TCA CA 3’ | 5’ TTC TGC AAG TGC ATC ATC GTT 3’ | NM_031168.2 |
| **MCP-1** | 5’ CAT CCA CGT GTT GGC TCA 3’ | 5’ GAT CAT CTT GCT GGT GAA TGA GT 3’ | NM_011333.3 |
| **Ly6C** | 5’ TCT TGT GGC CCT ACT GTG TG 3’ | 5’ AAA GAA AGG CAC TGA CGG GT 3’ | NM_010741 |
| **NeuN** | 5’ TGG CTG GAA GCT AAA CCC TG 3’ | 5’ CCC ACG GGA CTG GCA TTT TA 3’ | NM_001039167 |
| **NOS2** | 5’ TGG TGA AGG GAC TGA GCT GT 3’ | 5’ TGA GAA CAG CAC AAG GGG TTT 3’ | NM_010927.4 |
| **TNFα** | 5’ ATG GCC TCC CTC TCA TCA GT 3’ | 5’ TTT GCT ACG ACG TGG GCT AC 3’ | NM_013693.3 |
| **SmRNA** | 5’ CGG GAC AAG AAG GTG GAA G 3’ | 5’ AGT CTG CAG TGA GTC GAA GAA A 3’ | WO2004.092414 |

**Abbreviations**: CB1, cannabinoid receptor type 1; CB2, cannabinoid receptor type 2; CD, cluster of differentiation; COX2, cyclooxygenase 2; GFAP, glial fibrillary acidic protein; GLAST, glutamate aspartate transporter; GPR, G-protein coupled receptor; Iba1, ionized calcium-binding adapter molecule 1; IL, interleukin; MCP1, monocyte chemoattractant protein 1; NeuN, neuronal-nuclei; NOS2, Nitric Oxide Synthase 2; TNFα, tumor necrosis factor α; SmRNA, standard messenger RNA.
