## Supplementary material for "Cannabinoid receptor type 2 expression in mouse brain: from mapping to regulation in microglia under inflammatory conditions": Table S2

**Table S2 – Multiple comparison of CB2, CB1, GPR55 and GPR18 transcript levels measured in 6 different brain regions and 3 mouse strains.** Normality of CB2, CB1, GPR55 and GPR18 transcript level data distribution was tested using Shapiro–Wilk test and quantile–quantile plots ("Normality?" lines). Normal data were analyzed with Tukey's post-hoc test for multiple comparisons following one-way ANOVA. P-values of post-hoc tests are presented in the table.

| **CB2** | | | | | | |
|---|---|---|---|---|---|---|
| Brain region | **OB** | **NCX** | **HI** | **Hθ** | **CRB** | **BS** |
| Normality ? | Yes | Yes | No | Yes | No | Yes |
| Test | ANOVA 1 | ANOVA 1 | Kruskal-Wallis | ANOVA 1 | Kruskal-Wallis | ANOVA 1 |
| Swiss vs. Balb/c | 0.3032 | 0.9566 | 0.4081 | 0.9685 | >0.9999 | 0.7607 |
| Swiss vs. C57Bl/6 | >0.9999 | 0.7099 | >0.9999 | 0.7462 | >0.9999 | 0.9233 |
| Balb/c vs. C57Bl/6 | 0.0512 | 0.5501 | 0.6991 | 0.8714 | >0.9999 | 0.546 |
| **CB1** | | | | | | |
| Brain region | **OB** | **NCX** | **HI** | **Hθ** | **CRB** | **BS** |
| Normality ? | No | Yes | Yes | Yes | No | Yes |
| Test | Kruskal-Wallis | ANOVA 1 | ANOVA 1 | ANOVA 1 | Kruskal-Wallis | ANOVA 1 |
| Swiss vs. Balb/c | >0.9999 | >0.9999 | 0.277 | 0.4667 | >0.9999 | 0.8558 |
| Swiss vs. C57Bl/6 | 0.6991 | 0.3032 | 0.1226 | 0.9743 | 0.4081 | 0.3995 |
| Balb/c vs. C57Bl/6 | >0.9999 | 0.0512 | 0.8007 | 0.5819 | 0.6991 | 0.6831 |
| **GPR55** | | | | | | |
| Brain region | **OB** | **NCX** | **HI** | **Hθ** | **CRB** | **BS** |
| Normality ? | Yes | Yes | Yes | Yes | Yes | Yes |
| Test | ANOVA 1 | ANOVA 1 | ANOVA 1 | ANOVA 1 | ANOVA 1 | ANOVA 1 |
| Swiss vs. Balb/c | 0.8568 | 0.4659 | 0.8589 | 0.7817 | 0.935 | 0.5084 |
| Swiss vs. C57Bl/6 | 0.6712 | 0.5249 | 0.9994 | 0.6037 | 0.4309 | 0.4513 |
| Balb/c vs. C57Bl/6 | 0.9376 | 0.9928 | 0.8434 | 0.9466 | 0.2865 | 0.993 |
| **GPR18** | | | | | | |
| Brain region | **OB** | **NCX** | **HI** | **Hθ** | **CRB** | **BS** |
| Normality ? | Yes | Yes | Yes | Yes | Yes | Yes |
| Test | ANOVA 1 | ANOVA 1 | ANOVA 1 | ANOVA 1 | ANOVA 1 | ANOVA 1 |
| Swiss vs. Balb/c | 0.1211 | 0.8902 | >0.9999 | 0.7098 | 0.8291 | 0.9294 |
| Swiss vs. C57Bl/6 | 0.3208 | 0.4081 | 0.076 | 0.9991 | 0.8896 | 0.8808 |
| Balb/c vs. C57Bl/6 | 0.7287 | **0.0338** | 0.2209 | 0.6875 | 0.9911 | 0.9923 |

**Abbreviations**: BS, brain stem; CB, cannabinoid receptor; CRB, cerebellum; GPR, G protein coupled receptor; HI, hippocampus; Hθ, hypothalamus; NCX, neocortex; OB, olfactory bulb.
